# Bayesian stroke modeling details sex biases in the white matter substrates of aphasia

**DOI:** 10.1101/2022.01.18.474989

**Authors:** Julius M. Kernbach, Gesa Hartwigsen, Jae-Sung Lim, Hee-Joon Bae, Kyung-Ho Yu, Gottfried Schlaug, Anna Bonkhoff, Natalia S. Rost, Danilo Bzdok

## Abstract

Ischemic cerebrovascular events often lead to aphasia. Previous work provided hints that such strokes may affect women and men in distinct ways. Women tend to suffer strokes with more disabling language impairment, even if the lesion size is comparable to men. In 1,401 patients, we isolated data-led representations of anatomical lesion patterns and hand-tailored a Bayesian analytical solution to carefully model the degree of sex divergence in predicting language outcomes ∼3 months after stroke. We located lesion-outcome effects in the left-dominant language network that highlight the ventral pathway as a core lesion focus across different tests of language performance. We provide newly detailed evidence for sex-specific brain-behavior associations in the domain-general networks associated with cortico-subcortical pathways, with unique contributions of the fornix in women and cingular fiber bundles in men. Our collective findings suggest diverging white matter substrates in how stroke causes language deficits in women and men. Clinically acknowledging such sex disparities has the potential to improve personalized treatment for stroke patients worldwide.

## Introduction

Stroke is the leading cause of long-term disability worldwide. By 2047, the prevalence of stroke is estimated to increase by 27%, with women being disproportionately affected (Bushnell et al., 2014; Wafa et al., 2020). Estimations suggest a ∼55,000 increase of stroke events in women each year (Bushnell et al., 2014; Giroud et al., 2017). The anticipated rise in stroke prevalence will put a strain on the economic burden of our health systems. The annual stroke-related social and health costs have reached >60€ billion in Europe alone (Luengo-Fernandez et al., 2020; Olesen et al., 2012). Post-stroke aphasia places an overwhelming burden on many of the >25 million stroke survivors worldwide (Engelter et al., 2006; Feigin et al., 2017). For affected individuals, the sequelae of stroke can prompt drastic life changes, with severe long-term impairments (Feigin et al., 2014; Pedersen et al., 2004). Aphasia severely impacts everyday communication and social interactions by affecting language production and comprehension. Aphasia is associated with an increased mortality rate (Laska et al., 2001), longer hospitalization, poorer functional recovery, and reduced overall probability of returning to work (Gialanella & Prometti, 2009).

As an underappreciated fact, women typically suffer strokes with more severe consequences and language impairment even if the lesion size is comparable to men (Hier et al., 1994; Silva et al., 2010). Yet, the majority of stroke lesion studies have been underpowered to detect robust sex disparities given the surfeit of small-sample investigations (Button et al., 2013; Nord et al., 2017). For that reason, placing special emphasis on sex effects requires well-powered patient samples, especially given that many sex-related neural correlates are modest in effect size (Joel et al., 2015). The present investigation in a large patient cohort aims at a direct probabilistic assessment of the subtle sex effects in how the topography of white matter tissue damage leads to differences in language impairments. Such insights on sex-specific lesion-outcome associations are needed to improve personalized treatment prospects in light of the increased burden of aphasia in women.

Aphasia is a frequent consequence of embolic occlusion of the middle cerebral artery. However, brain lesion distributions are highly variable in the preserved and impaired language capacity. In attempting to harmonize the heterogeneous clinical manifestation of aphasia, decades of research efforts have been devoted to determining the complex neural basis of language processing. Today, there is growing consensus that the human language faculty is supported by distributed large-scale networks. Numerous studies have shed light on the anatomical white matter connections at play. In particular, the arcuate fasciculus (AF) has been at the center of attention for decades (Catani & Mesulam, 2008; Dick & Tremblay, 2012). Early anatomical dissection outlined its anatomical trajectory, which suggested a direct fiber connection of Wernicke’s area in the left temporal lobe to Broca’s area in the left frontal lobe (Dejerine et al., 1895; Geschwind, 1970). Applications of diffusion tensor imaging techniques promoted the idea of a further division of the AF into three segments. One direct temporo-frontal segment and two lateral segments have been proposed to compose the overall AF. The lateral tract subsegments implement an indirect fronto-parietal and temporal-parietal pathway (anterior and posterior segment, respectively) (Catani et al., 2005; Catani & Thiebaut de Schotten, 2008). The AF may provide an anchor for the canonical perisylvian network and has been placed at the core of the neuroarchitecture of the uniquely human language abilities (Rilling et al., 2008).

Attempting to marry its structural and functional bases, researchers have integrated the core perisylvian pathway into different functional-anatomical modules. One popular idea is the dual-stream concept of language (G. Hickok & Poeppel, 2000; Gregory Hickok, 2009; Gregory Hickok & Poeppel, 2004, 2007; Rauschecker & Tian, 2000; Saur et al., 2008). This view puts forward a ventral stream for mapping sound-to-meaning and a dorsal stream comprising the AF and superior longitudinal fascicle for mapping sound-to-motor representations. The ventral pathway includes the uncinate fasciculus (UF), the inferior fronto-occipital fasciculus (IFOF), and inferior longitudinal fasciculus (ILF) that together assist in processing speech signals for language comprehension (Catani et al., 2003; Catani & Mesulam, 2008; Papagno et al., 2011; Saur et al., 2008). The dorsal stream is thought to be left-dominant, while the ventral stream is believed to be more bilaterally organized (Catani & Mesulam, 2008; Eichert et al., 2019; Gregory Hickok & Poeppel, 2007). However, there is strong interindividual variability in the degree of hemispheric dominance in general, and accumulating evidence points towards aspects of sex-specific differentiation in particular. For example, more pronounced left lateralization in the long segment of the AF was found in men (Thiebaut de Schotten et al., 2011). Sex differences have also been reported for lateralization of functional activation during carefully designed language tasks (Shaywitz et al., 1995).

Clinically, ischemic stroke generally affects male and female patients differently in terms of incidence, severity, survival, and potential for recovery (Bonkhoff, Karch, Weber, Wellmann, & Berger, 2021; Bushnell et al., 2014; Dehlendorff et al., 2015). In fact, women generally suffer strokes with more severe neuropsychological consequences compared to those of men (Bonkhoff, Schirmer, Bretzner, Hong, et al., 2021; Silva et al., 2010). Similar lesion volume effects were noted in cases of aphasia, in which smaller ischemic stroke lesions were capable of producing aphasia in women than men (Hier et al., 1994). The reasons behind sex idiosyncrasies in functional outcomes after stroke remain elusive. Differences are often explained by poorer pre-stroke function, more comorbidities such as depression, and social isolation at an older age (see (Reeves et al., 2008)). Yet, adjustment for these factors does not adequately explain the observed discrepancies in stroke outcomes between men and women (Di Carlo et al., 2003; Holroyd-Leduc et al., 2000).

Despite the apparent differences in stroke outcome between women and men, sex is often treated as a nuisance variable in the overwhelming majority of stroke lesion studies (Baldo et al., 2013; Mirman et al., 2015; Wu et al., 2015). This established practice blinds researchers to detect any potential neuroanatomical effects that depend on sex. Compounding these challenges that hold barriers to understanding sex-specific brain lesion effects, lesion-symptom mapping studies often constrained the analyses to the predominantly affected hemisphere. That is, as a general trend, the left hemisphere is specifically investigated in language impairment and the right hemisphere in neglect (Baldo et al., 2013; Bates et al., 2003; Bonilha et al., 2017; Smith et al., 2013; Wu et al., 2015). To overcome several existing roadblocks, the present work takes advantage of Bayesian principles as a formal device to quantify degrees of sex divergences. This clean methodology enabled sex-aware analysis in 1,401 ischemic stroke patients, covering major white matter pathways in both hemispheres. Understanding how lesion location, structural language lateralization, and their clinical outcome diverge in aphasia between female and male populations can finess personalized rehabilitation and treatment.

## Results

We go new ways to chart sex disparities in one of the largest existing stroke cohorts with rigorous neuropsychological profiling. We espoused probabilistic Bayesian principles for direct uncertainty quantification of nuanced sex effects on language performance attributable to specific white matter lesions. First, we extracted a low-dimensional embedding of the voxel-granularity MRI-lesion distributions using unsupervised pattern discovery. Second, the patient-specific expressions of the derived lesion configurations provided the basis for a Bayesian model to predict stroke outcomes with its full probabilistic differentiation between women and men.

### Patient cohort: characteristics and behavioral outcome measures

Our inclusion criteria were met by 1,401 patients with ischemic stroke who were recruited as a prospective multi-center cohort from 2007-2018 (**Table 1** for details). The mean age was 67.58 (standard deviation 11.59), with 42% women. The mean age at admission was significantly higher in females than males (t=8.9, p<0.001, cf. Table 1). Patients had an average of 9.28 (5.92) years of education, with women amounting to fewer years on average compared to those of men (t=-17.27, p<0.001). Based on the IQCODE score, females’ cognitive performance before the ischemic event was significantly worse compared to males (t=3.91, p<0.001). Stroke severity as reflected in the clinical impairment was significantly higher in females across nine cognitive tests (K-MMSE, BN, SF, PF, RCFT, RCFT delayed recall, digital symbol coding, TMT-A, and -B). Out of 15 objects to be recognized in the BN (Kaplan et al., 1983), patients correctly named 9.92 (3.59) on average, with females scoring worse than males (t=-11.90, p<0.001). Females also performed worse in semantic and phonemic fluency (t=-7.12 and -7.97, both p<0.001). In sum, we observed more severe stroke-related deficits and worse premorbid cognitive performance in female patients of our cohort.

**Table 1.** Characteristics of the patient cohort. Variables depicted as mean (standard deviation). Sex differences were tested using independent t-test (t, p-value), or in case of unequal variances using Welch’s t-test (t, p-value).

|  | Cohort | Women | Men | Sex differences |
| --- | --- | --- | --- | --- |
| Age | 67.58 (11.59) | 70.73 (11.15) | 65.29 (11.38) | 8.9, <0.001 |
| Lesion volume | 17.08 (43.33) | 15.05 (44.16) | 18.57 (42.70) | -1.5, 0.1337 |
| IQCODE | 3.36 (0.46) | 3.42 (0.49) | 3.32 (0.44) | 3.91, <0.001 |
| Years of education | 9.28 (5.20) | 6.68 (4.98) | 11.16 (4.51) | -17.27, <0.001 |
| K-MMSE | 23.73 (5.92) | 21.93 (6.34) | 25.03 (5.23) | -9.98, <0.001 |
| Boston naming (BN) | 9.92 (3.59) | 8.63 (3.54) | 10.86 (3.32) | -11.90, <0.001 |
| Semantic fluency (SF) | 11.51 (5.11) | 10.39 (4.89) | 12.33 (5.12) | -7.12, <0.001 |
| Phonemic fluency (PF) | 17.37 (10.78) | 14.76 (10.16) | 19.27 (10.82) | -7.97, <0.001 |
| SVLT immediate recall | 14.82 (6.01) | 14.71 (6.45) | 14.90 (5.67) | -0.57, 0.5 |
| SVLT delayed recall | 3.67 (2.92) | 3.73 (3.04) | 3.64 (2.82) | 0.54, 0.5 |
| SVLT recognition | 18.16 (4.14) | 17.96 (4.14) | 18.31 (4.14) | -1.55, 0.1 |
| RCFT delayed recall | 11.33 (5.83) | 9.49 (5.38) | 12.66 (5.78) | -10.42, <0.001 |
| RCFT | 25.74 (8.68) | 23.09 (9.35) | 27.67 (7.60) | -10.09, <0.001 |
| Digital symbol coding | 35.67 (21.73) | 29.35 (20.90) | 40.26 (21.17) | -9.56, <0.001 |
| TMT-A | 42.53 (36.52) | 52.81 (46.70) | 35.11 (24.36) | 9.21, <0.001 |
| TMT-B | 95.58 (76.56) | 117.80 (85.56) | 79.28 (64.41) | 9.61, <0.001 |

### Topography of ischemic stroke lesion patterns

Excellent whole-brain coverage was attested by the voxel-wise summary of lesion overlap across 1,401 patients (**Figure 1A**). The lesion overlap predominantly corresponded to the vascular territory of the left and right middle cerebral artery. The majority of lesions were located in the deep central white matter in both hemispheres. Most tissue lesions were concentrated in the thalamus and basal ganglia. The average lesion volume was 17.08 ml (14.81 - 19.36) and did not significantly differ between female and male patients within our cohort (women: 15.05 ml (11.47 - 18.63), men: 18.57 ml (15.63 - 21.51), *p*=0.13; **Table 1**). We did not find significant differences between females and males regarding their stroke topography (**Figure 1B-C,** summarized separately).

**Figure 1.**
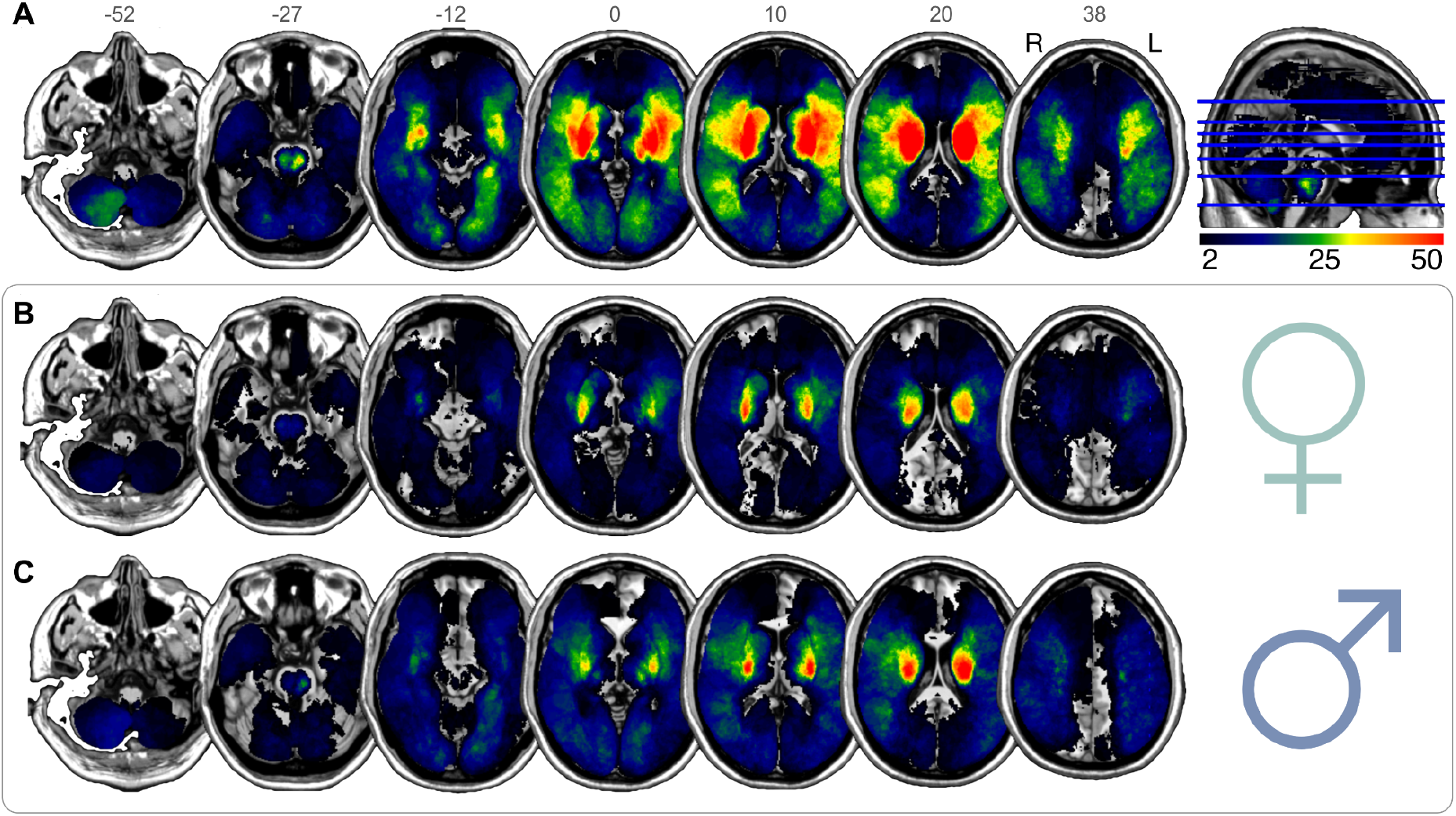
Spatial topography of ischemic stroke lesions summarized from 1,401 patients. We confirm excellent whole-brain coverage, without appreciable hemispheric differences in the voxel-level stroke topography for the whole cohort (**A**), or separately either for female patients (**B**) or for male patients (**C**). The overall lesion volume did not significantly differ between female and male patients within our cohort (two-sided t-test, *p*=0.13, cf. Table 1). The topographical distributions of tissue damage caused by ischemic stroke are shown in MNI reference space with z-coordinates indicated above each brain slice. Lesions predominantly overlapped in areas that correspond to the vascular territory of the left and right middle cerebral artery. The majority of lesions were localized in subcortical zones, which entails a disconnection of deep white matter tracts. Radiological view; R/L: right/left.

### Anatomy of the extracted stroke lesion atoms

We first extracted hidden topographical archetypes from the high-dimensional lesion distribution in white matter at voxel resolution. Using non-negative matrix factorization (NNMF; (D. D. Lee & Seung, 1999)) for unsupervised pattern discovery, we uncovered recurring spatial configurations of concurrent tissue damage that are consistently apparent across >1,000 stroke cases - which we call *lesion atoms* **(Figure 2)**. Disentanglement of this sum-of-parts representation by means of machine learning provides an important advantage for the purpose of the present study. The non-negativity constraint of the applied algorithm enabled neurobiological meaningful and direct interpretable lesion patterns. In contrast, alternative dimensionality reduction tools such as principal component analysis would hurt an intuitive interpretation of the lesion atom effects. This is because alternative tools recover patterns through incomprehensible positive and negative cancellations of the separate lesion configurations. As a descriptive post-hoc analysis, each lesion atom was assessed for hemispheric dominance of the implicated tissue lesion distribution by an index of lateralization (**Figure 2C**). For most factors, there was a unique left-hemispheric lesion constellation in correspondence with a similar right-hemispheric homolog (**Figure 2A box,** factors 3 and 9, or 2 and 5). Overall, the spatial distribution of the derived lesion atoms emerged to be directly interpretable and biologically plausible with components underlying distinct territories of arterial supply (**Figure 2**).

**Figure 2.**
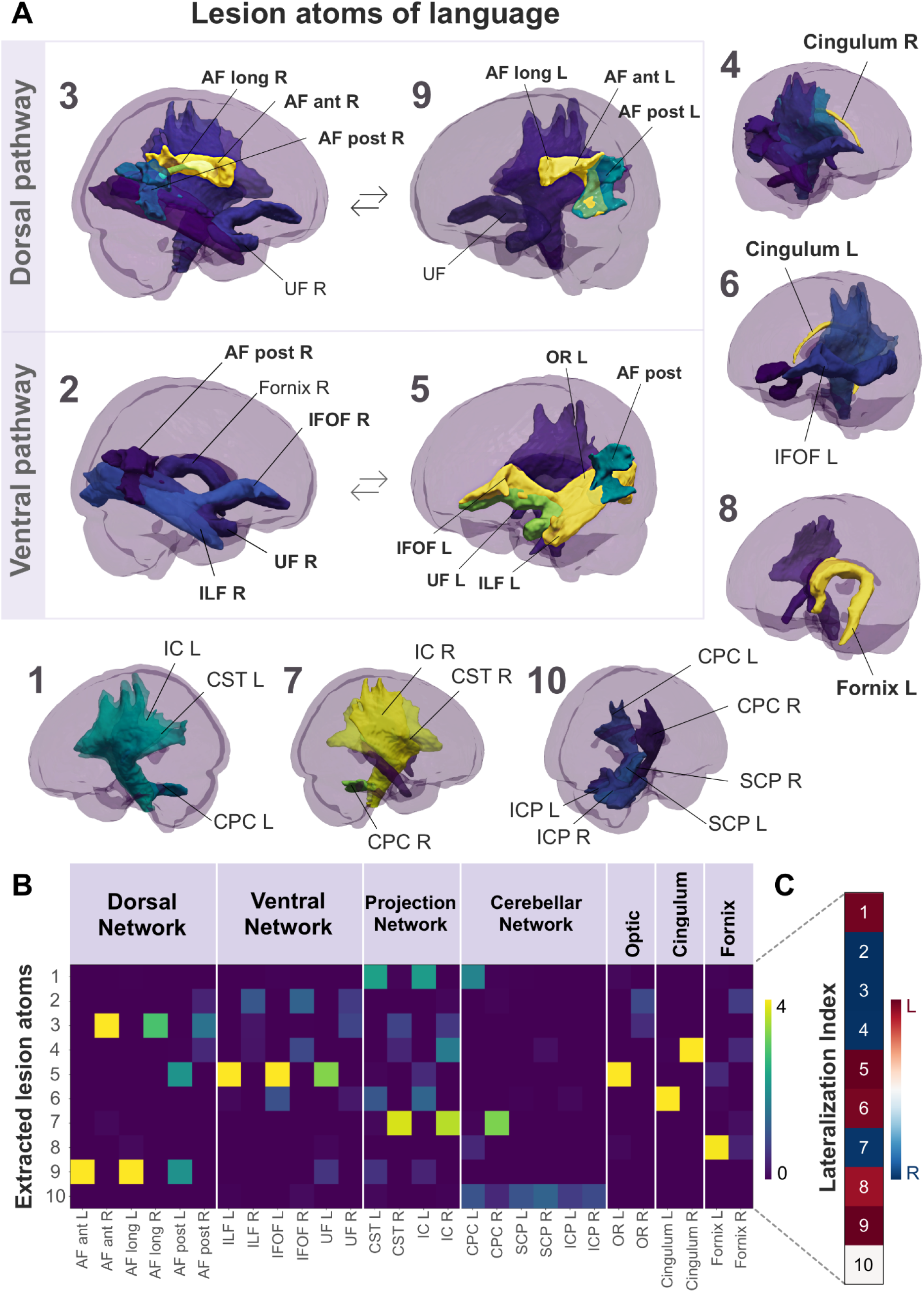
Multivariate latent-factor discovery deconvolves unique stroke lesion atoms. In 1,401 neurological patients, we used an unsupervised machine learning algorithm to perform a data-driven discovery of white matter lesion patterns. This sum-of-parts approach (non-negative matrix factorization, NNMF) enabled the identification of archetypical white-matter lesion configurations, *lesion atoms,* distilled from high-resolution brain scans with a total >1,7 million 1mm^3^ lesion voxels across patients. The anatomical distribution of the derived lesion atoms corresponded to directly interpretable and biologically plausible lesion patterns that are reminiscent of well-known functional systems (**B**). Each atom of coherent white matter lesion was assessed for hemispheric asymmetry (**C**) based on a lateralization index (LI) (Ito & Liew, 2016). This index is computed from the weights of the latent factor loadings by LI = (L-R)/(L+R). Hemispheric asymmetry effects are indicated by left- (Red) or right-lateralization (Blue). Most factors showed a left-hemispheric lesion configuration in correspondence with a similar right-hemispheric homolog. Four factors were predominantly associated with the language-related pathways (**A**, Box): the dorsal perisylvian circuitry (Factor 3 and 9; R and L, respectively) and the ventral pathway (Factor 2 and 5; R and L, respectively). Notably, the core perisylvian pathway shows significant leftward asymmetry in factor 9 with the accentuated influence of the long and anterior AF. Instead, the right homolog in factor 3 predominantly relies on the anterior segment of the AF. Motor function (**A**, bottom row) was most strongly expressed in factors 1 (L) and 7 (R), and 10 (bilateral cerebellar network). Primarily limbic factors 4 and 6 were dominated by the influence of the left and right cingulum. Factor 8 was uniquely left-lateralized and captured the main effect of the fornix (**A**, right column). As such, even subtle differences in lateralization align with the current neuroanatomical knowledge of the language pathways (Thiebaut de Schotten et al., 2011). Abbreviations: Arcuate fasciculus: AF. Inferior longitudinal fasciculus: ILF. Uncinate fasciculus: UF. Inferior fronto-occipital fasciculus: IFOF. Cortico-spinal tract: CST. Internal capsule: IC. Cortico-pontine-cerebellar tract: CPC. Inferior and superior cerebellar peduncle: ICP, SCP. Optic radiation: OR. R/L: right/left.

Four lesion atoms of stroke-induced tissue damage were mapped explicitly to anatomical tracts implicated in the dorsal and ventral language pathways (**Figure 2A**). Generally, the left-lateralized language atoms showed stronger lesion effects than their right-lateralized homologs. Factor 9 primarily covered white-matter lesions of the core perisylvian pathway with a focus on the left hemisphere. These lesion effects predominantly rested on the AF. The level of relevance within the subsegments of the AF followed an anterior-to-posterior gradient. Here, the anterior and long segments showed higher lesion importance than the posterior segment. The core perisylvian right-hemispheric homolog was captured by factor 3. The organization of the AF’s different segments again showed an anterior-to-posterior transition. Both factors implicated the UF pathway to a lesser extent. Factor 5 resembled the left-lateralized ventral pathways, which included the ILF, the IFOF, and the UF. These lesion atom effects further implicated damage in the left optic radiation (OR), and to a lesser degree, the posterior segment of the AF. The right-hemispheric homolog to factor 5 was identified as factor 2. Right-hemispheric (Factor 2) ventral pathways were less pronounced regarding their overall lesion importance compared to the left.

Three lesion atoms resembled damage to the brain’s motor circuits (**Figure 2A** bottom row). The pattern topology of lesion factor 1 included the left-lateralized motor projections of the cortico-spinal tract (CST), internal capsule (IC), and cortico-pontine-cerebellar tract (CPC). Similarly, factor 7 captured lesions in the mirrored motor projections of the right hemisphere. Factor 10 was the only factor with no hemispheric dominance and captured bilateral cerebellar connections, including the CPC, the inferior and superior cerebellar peduncle (ICP, SCP). Three further lesion atoms captured the prominent influence of the limbic system. The right cingular fibers dominated the tissue damage configuration of factor 4, next to a lesser influence of the posterior segment of the AF, IFOF, and IC. Conversely, the lesion constellation of factor 6 captured the left-hemispheric cingular influence. Lastly, factor 8 was dominantly shaped by the influence of the left fornix. Notably, only factor 8 did not have a homologous pattern on the right hemisphere.

By bringing to bear an algorithmic approach from machine learning, we deconvolved unique lesion atoms as low-dimensional representations of stroke configurations from 1,401 patients. Out of ten lesion atoms, four atoms specifically resembled the ventral and dorsal core language pathways on the left and homologous right hemisphere. Three resembled motor pathways, and three further lesion atoms predominantly coincided with limbic pathways in the fornix and cingular fiber pathways.

### Lesion atoms track interindividual differences in cognitive outcomes

To chart the clinical relevance of each lesion atom, we computed Pearson’s correlations between patient-specific expressions of a given lesion atom and a cognitive test, one for each available neuropsychological assessment (**Figure 3**). Based on the ensuing (absolute) factor-outcome association strengths (**Figure 3A**), we could rank the lesion atoms across all outcome measures of our cohort with unusual deep phenotyping. As a consequence of this bottom-up approach, factors 5 (Ventral pathway R), 8 (Fornix L), and 9 (Dorsal pathway L) ranked as the three most clinically relevant lesion atoms given our functional domains of cognitive outcomes. The limbic factors 4 and 6 reflected the right and left influence of the cingular fiber bundle and ranked fourth and fifth. Except for the TMT, factors 5, 8, and 9 showed pronounced negative associations across most cognitive scores (**Figure 3B**). Regarding the language outcomes, damage in the left-hemispheric lesion atoms of the ventral and dorsal pathways (Factors 5 and 9, respectively) caused a significant degree of deterioration in Boston naming (BN), semantic fluency (SF), and phonemic fluency (PF). However, the association with their respective right-hemispheric homologs diverged: Notably, the right hemispheric homolog of the core perisylvian network (Factor 3) showed a slightly negative association with BN, but otherwise positive associations for SF and PF. In contrast, the right-hemispheric homolog of the ventral pathway (Factor 2) was negatively associated with BN and SF, with close links to deteriorated naming and speech fluency.

**Figure 3.**
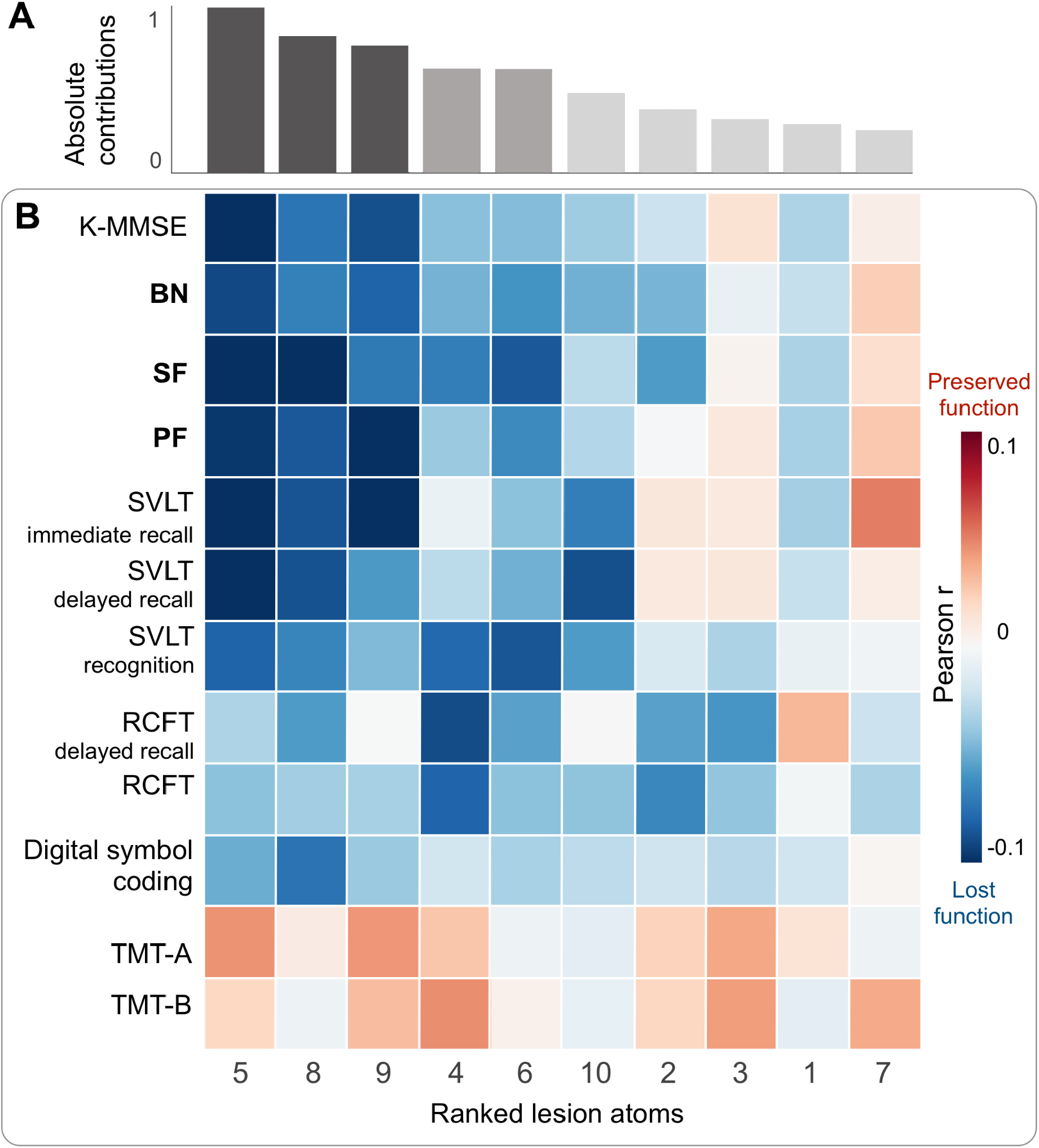
Stroke lesion atoms are ranked from strongest to weakest associations with clinical outcomes. We assessed the overall clinical relevance of each lesion atom using Pearson’s correlation between subject-specific lesion atom expression and a cognitive test across outcome measures. **A**) Based on the summed absolute contributions, factors 5 (Ventral pathway L), 8 (Fornix L), and 9 (Dorsal pathway L) ranked as the three most relevant lesion atoms. Following the top three lesion atoms, the limbic factors 4 and 6 reflect the right and left influence of the cingular fiber bundles. **B**) All top five lesion atoms showed pronounced negative associations with cognitive impairment across most neuropsychological assessments. Regarding language, damage in the left-hemispheric lesion atoms (Factors 5 and 9) of the ventral and dorsal pathway cause a significant degree of deterioration in the Boston naming test (BN), semantic and phonemic fluency (SF, PF). However, the association with their respective right-hemispheric homologs diverged: the right-hemispheric homolog of the dorsal core perisylvian network (Factor 3) showed slight negative correlation in BN, but otherwise positive correlations in SF and PF. In contrast, the right-hemispheric homolog of the ventral pathway (Factor 2) is associated with a considerable deterioration in BN and SF. Abbreviations: Korean-Mini Mental State Examination: K-MMSE. Boston Naming: BN. Semantic fluency: SF. Phonemic fluency: PF. Seoul-Verbal Learning: SVL. Rey Complex Figure Test: RCFT. Korean-Trail Making Test Version A/B: TMT A/B. R: right, L: left.

We thus ranked the uncovered lesion atoms based on their clinical sequelae. Specifically for language outcomes, we observed diverging responses in the left and right homologs of the ventral and dorsal pathways: for the dorsal perisylvian pathway, severe deterioration was found for left-hemispheric damage (Factor 9) and less for right-hemispheric damage (Factor 3). In contrast, both damage to the left (Factor 5) and right (Factor 2) ventral pathways especially inflicted impairment in both naming and speech fluency.

### Sex divergences in lesion atom expressions and prediction of language outcome

Next, we turned to the direct quantification of degrees of sex differentiation in how the emerged lesion patterns predict language outcome. Based on the patient-specific constellations of lesion pattern expressions, we estimated a fully probabilistic, generative model to explain language outcome at ∼3 months after stroke. In particular, our approach was able to obtain full Bayesian posteriors over the differentiation between females and males. In the present analysis, we focused on the quantitative analyses of the language outcomes of naming ability (BN) and speech fluency (SF and PF; other scores c.f. **Supplement Fig. 1-12**). Leveraging the Bayesian modeling framework, we provided an honest quantification of a) the general effects of the derived lesion atoms on cognitive outcome after stroke and b) the degree of sex divergence in how the lesion atoms achieve the prediction of post-stroke language outcome. All analyses were adjusted for variation that could be explained by age, age^2^, sex, education in years, pre-morbid cognitive performance, and total lesion volume. We also included sex as a covariate in the model to explicitly account for a-priori differences between female and male stroke outcomes (Bonkhoff, Schirmer, Bretzner, Hong, et al., 2021). For example, worse language outcomes in females could be linked to an underlying cause independent of the lesion distribution, such as a decreased likelihood of receiving acute interventions. In this scenario, the effect would be expressed in the Bayesian posterior distribution of the sex covariate, but not in that of the female-stratified lesion atom.

Among our target language capacities, we first examined interindividual differences in naming ability as captured by the BN test. The posterior predictive checks confirmed an appropriately fitted approximation of the data variation that underlies how lesion atom expressions explain cognitive performance outcomes (R^2^=0.51, **Figure 4A**). We then inspected the role of each lesion atom in forecasting future naming performance in BN at ∼3 months after stroke, separately for women and men (**Figure 5A**). Both sexes displayed worse naming outcomes with increasing damage to the left-lateralized ventral pathway (Factor 5), its right-hemispheric homolog (Factor 2), and the right cingular fiber pathway (Factor 4). However, the inferred posterior parameter distributions also attested specific divergences between females and males regarding lesion-outcome associations: the right-hemispheric homolog of the core perisylvian network (Factor 3) and the left fornix (Factor 8). Here, damage in the lesion pattern configurations apparent in men was predictive of worse language outcomes. In contrast, stronger impairment in females was further explained by damage in the left core perisylvian network (Factor 9). Tissue damage in the motor projections of the right hemisphere (Factor 7) and the left cingular fiber pathway (Factor 6) explained more pronounced deficits in naming performance in women than men. We thus identified common and diverging effects across the subject expressions of lesion atoms to predict naming capacity between women and men.

**Figure 4.**
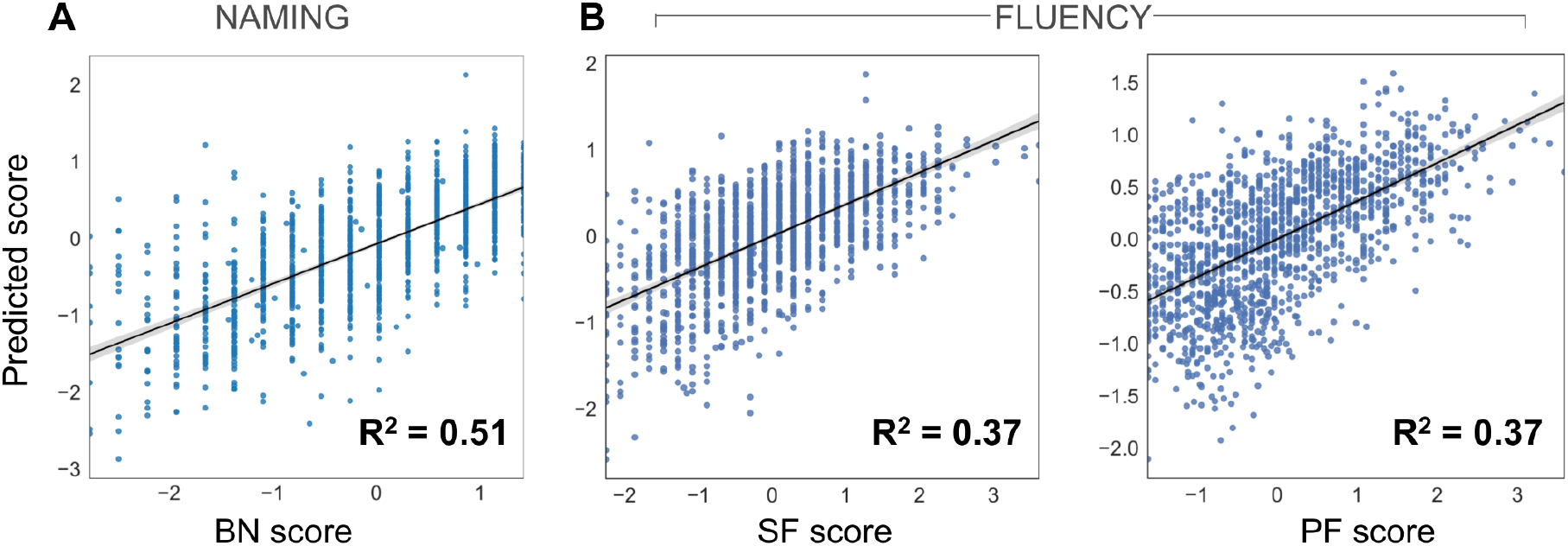
Performance of the inferred Bayesian analytical solution in predicting naming and speech fluency. We confirmed a well-fitted approximation of the underlying distribution as performed for model checking in previous research (Bonkhoff, Lim, Bae, Weaver, Kuijf, et al., 2021; Kiesow et al., 2021). The posterior predictive checks are shown for the Bayesian models that were estimated to predict inter-individual difference in (z-scored) Boston naming (**A**, BN), semantic and phonemic speech fluency (**B**, SF and PF). These model-based simulations of new data were then compared to the actually observed data to compute the overall explained variance (coefficient of determination, R^2^). This practical check of model-based outcome predictions is a well-recognized approximation to external validation given the actual data at hand (Kruschke, 2014). The full model for the BN outcome explained a total variance of R^2^=0.51, and the Bayesian models for the PF and SF explained R^2^=0.37, respectively. Considering our white matter results, the predictive performance for naming and speech fluency matches previous research on grey matter lesions due to stroke (Bonkhoff, Lim, Bae, Weaver, Kuijf, et al., 2021).

**Figure 5.**
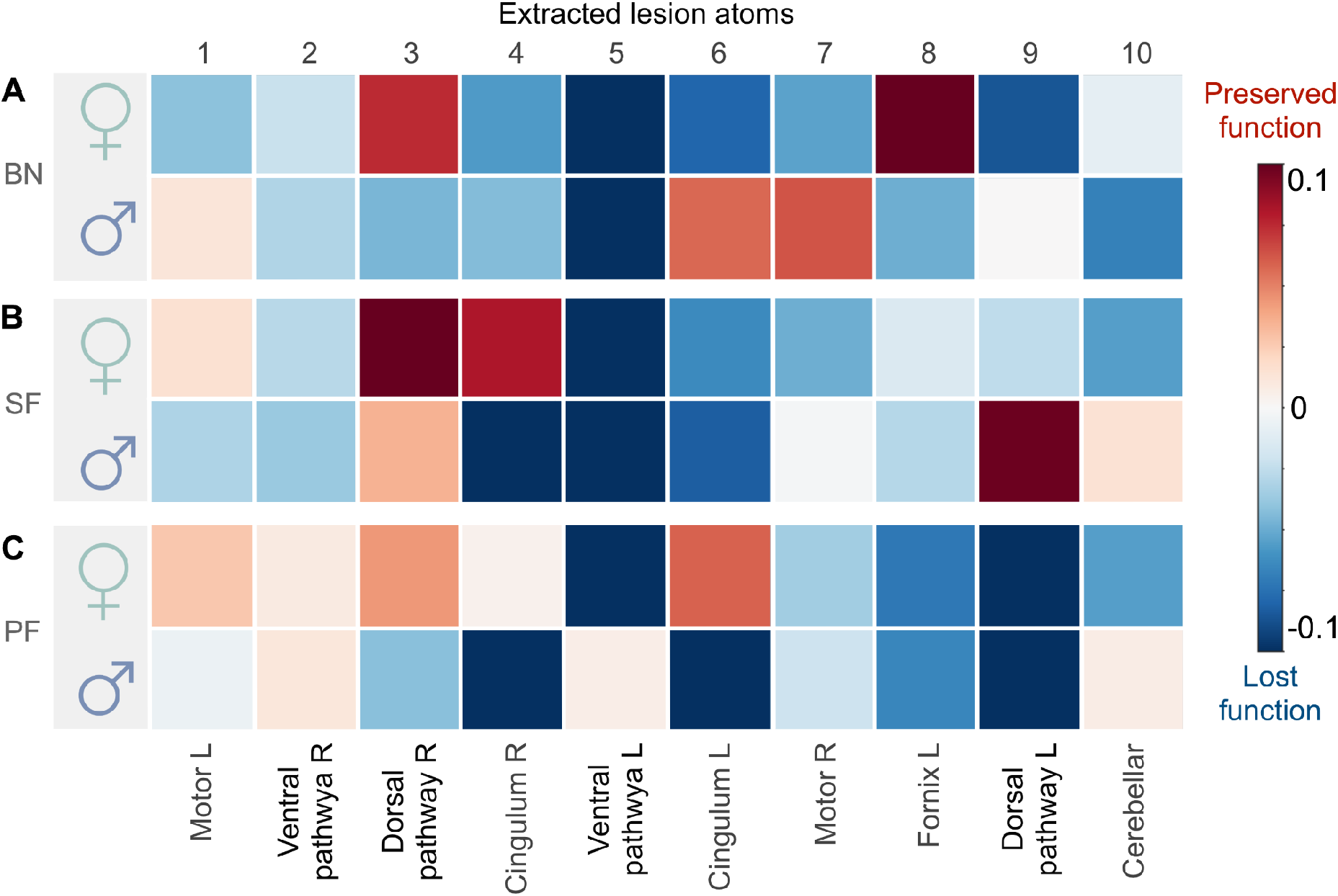
Sex bias becomes apparent from contributions of specific lesion atoms to explaining language outcomes. For each lesion atom, we show the predictive role in explaining language function separately for women and men. Shows the mean marginal Baysian posterior distribution of each lesion atom to explain moderate to severe language impairment. Across sexes, effects can be similar or diverging. **A**) For the Boston naming (BN), we observed driving loss of function common to both sexes in the left-lateralized ventral pathway (Factor 5), its right-hemispheric homolog (Factor 2), and the right cingulum (Factor 4). Diverging lesion effects between females and males arise in several atoms, including the right dorsal pathway, the left fornix, cingular tract, and the left and right motor projections (Factors 3, 8, 6, 7 and 1 respectively). **B**) Similarly for semantic fluency (SF), the main effect common to both sexes leading to worse performance was in the left ventral pathway (Factor 5) as well. Less influential were common effects in the right ventral pathway, the left cingulum and fornix (Factors 2, 6, 8). Analogous diverging lesion-outcome predictions in the right dorsal perisylvian circuit (Factor 3) were less pronounced than in BN, leading to moderate impairment for both sexes. But lesions in the left dorsal pathway lead to worse outcomes specifically for females. We further observed differences between women and men in the lesioned right cingulum (Factor 4). Here, males were predicted to have a more severe impairment. **C**) For phonemic fluency (PF), adverse outcome is primarily explained by the contributions of the left dorsal pathway (Factor 9) and left Fornix (Factor 8) in both sexes. We observed marked differences between sexes for the right-hemispheric dorsal pathway, left and right cingulum, with males performing worse than females (Factor 3, 4 and 6). In contrast, women showed significantly worse outcomes in the lesioned left ventral pathway (Factor 5).

Out of all extracted lesion atoms, distinct factors carried explanatory weight jointly for both sexes or uniquely for the female and male patients in our stroke population. The explanatory relevance for BN was indicated by the inferred marginal posterior parameter distributions (**Figure 6A**). Most relevant across both sexes, we identified a driving effect in the left-lateralized ventral pathway (Factor 5). Stroke-induced consequences were stronger in women (mean of the posterior distribution [PM] = -0.251, highest posterior density interval [HPDI] of the posterior distribution covering 80% certainty = -0.410 to -0.090) than in men (PM = -0.158, HPDI -0.267 to -0.040). We found further sex-diverging lesion-outcome effects. In women, tissue damage in the left fornix (Factor 8) predicted a more preserved outcome (Factor 8; PM = 0.138, HPDI 0.008 to 0.248). In males, we identified small but consistent effects: the right homolog of the ventral pathway contributed to the prediction of lost function (Factor 2; PM = -0.032, HPDI -0.063 to -0.009). Further, damage to the bilateral cerebellar network (Factor 10; PM = -0.080, HPDI -0.119 to -0.037) contributed to explaining naming impairments. Together, our BN findings witness a dominant lesion-outcome effect in the left ventral language pathway, consistently for both women and men. For women, the left fornix lesion was indicative of more moderate impairment of cognitive capacity ∼3 months after stroke. In contrast, males only showed modest diverging lesion pattern effects located in the homologous right ventral pathway and bilateral cerebellar connections.

**Figure 6.**
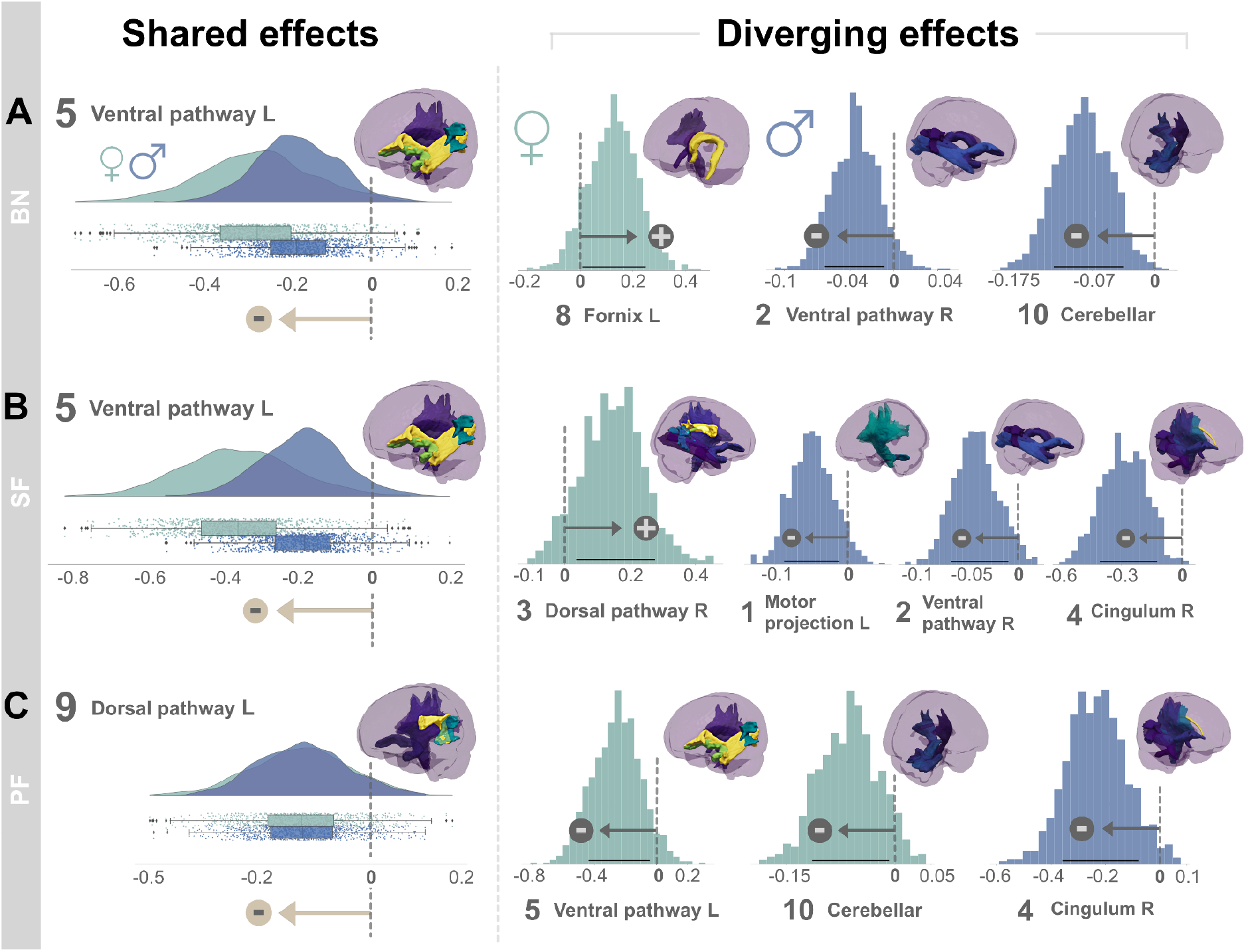
Shared and diverging lesion effects on language performance outcomes in female and male stroke populations. We formed full posterior-Bayes estimates to infer the explanatory relevance of each lesion atom for the prediction of language outcomes in female and male stroke patients. Key assessments of post-stroke language performance included the Boston Naming test (BN, **A**), semantic (SF, **B**) and phonemic fluency (PF, **C**). Lesion atoms were considered relevant if the highest probability density interval (HPDI) of the posterior distribution covering 80% certainty (black line) did not overlap with zero. We found lesion pattern effects that were common to both sexes (left side) and that diverged between women and men (right side). Across both sexes, lesion topographies pointed to an overarching structural organization reflecting the dual-stream concept of language (Gregory Hickok & Poeppel, 2004). Here, the ventral pathway was critically involved in naming and semantic fluency deficits, and the dorsal pathway in phonemic fluency deficits, congruently for women and men. More importantly, we provide novel evidence for sex-nuanced lesion-outcome effects. Women primarily rely on left-dominant circuits, including structural connections of the fornix, and a female-specific vulnerability to ventral lesion damage in phonemic fluency. In contrast, men draw from bilateral ventral and cerebellar connections, and potentially integrate more domain-general cognitive and executive functions via right-lateralized cingular fiber bundle to enable efficient language processing after stroke.

Examining how SF outcome differences are underpinned by white matter tissue damage, the posterior predictive check indicated an explained variance of R^2^=0.37 (**Figure 4B**). Considering the male versus female lesion-outcome associations, we identified strong effects for both sexes in the left ventral pathway (Factor 5; **Figure 5B**), followed by the left cingular fibers (Factor 6) and the right homolog of the ventral pathway (Factor 2). Damage captured by each of these three lesion atoms played contributing roles in lost SF function. Lesions affecting the right dorsal perisylvian circuit (Factor 3) explained a more moderate deficit for both sexes. Yet, in the SF function, too, we detected pronounced sex divergences. Men deteriorated more in SF tests as a result of tissue damage in the right cingular fibers (Factor 4). In contrast, white matter damage in the left core perisylvian network (Factor 9) added to difficulties with language fluency in women.

The lesion patterns predictive of SF differences resembled those explaining BN differences as evidenced by the corresponding marginal posterior distributions: our model attributed SF dysfunction in part to tissue damage in the left ventral pathway (Factor 5, **Figure 6B**) in both sexes. This lesion-outcome association was more pronounced in women (PM = -0.360, HPDI -0.561 to -0.186) than men (PM = -0.190, HPDI -0.345 to -0.069). Patient expressions of the lesion atom in the right-hemispheric perisylvian network were informative for predicting preserved function in women (PM = 0.157, HPDI 0.034 to 0.276) but not in men (**Figure 6 left**). Similar to the BN analysis, the right ventral pathway (Factor 2, PM = -0.040, HPDI -0.069 to -0.010) helped predict loss in SF function in males (**Figure 6 right**). The left-hemispheric motor projections damage (Factor 1, PM = -0.045, HPDI -0.088 to -0.012) helped predict SF dysfunction. However, we observed an even stronger lesion-outcome contribution by the right cingular fibers (Factor 4, PM = -0.268, HPDI -0.400 to -0.121) in men. Concluding our SF findings, we found a common lesion-outcome effect in the left ventral pathway. In women, tissue damage compromising the right perisylvian pathway added to the predictability of SF performance outcomes in our patients. Damage caused to the right cingulum assisted in the prediction of future SF loss in men beyond the damage caused to the left ventral pathway.

As the last examined facet of language capacity, we quantified how patterns of tissue damage underpin interindividual differences in PF performance ∼3 months after stroke. The full probabilistic model explained R^2^=0.37 of the variance in the outcome predictions based on the patient expressions of dedicated lesion atoms (**Figure 4B**). Common to both sexes, our model attributed lesion damage in the left core perisylvian circuit (Factor 9) to loss of PF function. We found marked divergences between sexes in factors 3-6 (**Figure 5**). Specifically, language dysfunction in females was due, to a larger extent, to damage to the left ventral pathway (Factor 5). In contrast, men showed more pronounced consequences for cognitive performance due to damage to the right and left cingulum (Factor 5 and 6) and the right perisylvian pathway (Factor 3).

Based on the inferred Bayesian posteriors (**Figure 6C**), the left dorsal perisylvian pathway showed considerable relevance in explaining differences in PF function, which was comparable in both sexes (Factor 9, females: PM = -0.151, HPDI -0.298 to -0.020; males: PM = -0.146, HPDI -0.273 to -0.029). However, women’s functional loss in PF was more explained by damage in the left ventral pathway (Factor 5, PM = -0.235, HPDI -0.437 to -0.043) compared to men. Additionally, lesions in the bilateral cerebellar network (Factor 10; PM = -0.058, HPDI -0.114 to -0.008) tracked performance decline in PF. Further, men showed a strong negative effect in the right cingular pathway (Factor 4, PM = -0.217, HPDI -0.359 to -0.077). Together, our lesion-outcome effects in PF highlighted the influence of the left-dominant perisylvian network common to both sexes. In women, the left ventral pathway added predictive value as to a loss of function in SF. In men, however, the influence of the right cingulum was further indicative of deterioration in PF.

Our sex-sensitive approach enabled us to tease apart shared and diverging lesion-outcome effects in female and male stroke populations. Across both sexes, the ventral pathway was critically involved in naming and semantic fluency deficits, while the dorsal pathway was implicated in phonemic fluency deficits. More generally, we provide novel evidence for sex-nuanced lesion-outcome effects. Concluding our findings across all language performance scores, women tend to rely on left-dominant circuits, including structural connections of the fornix. We further detailed a female-specific vulnerability to ventral lesion damage in phonemic fluency. In contrast, this facet of language performance in men tended to draw from bilateral ventral and cerebellar connections. Damage caused to the right cingular fiber connections assisted in the prediction of future SF loss in men beyond the damage caused to language-specific pathways.

## Discussion

Language capacity is uniquely evolved in humans and acts as a pacemaker in our societies. There is now an increasing realization that brain tissue damage caused by stroke entails partly distinct consequences on language performance in women and men. We hence designed a fully probabilistic framework as a clean analytical solution to modeling the apparent degrees of sex disparity. First, we extracted neurobiologically meaningful lesion patterns from 1,401 stroke patients in a data-led fashion. Second, we custom-tailored a bespoke Bayesian strategy to partition interindividual variation in caused aphasia outcomes into their lesion pattern contributions. In so doing, we offer honest uncertainty quantification of sex-biased effects in clinical language outcomes after stroke in one of the largest stroke cohorts with rarely available deep phenotyping.

In particular, (1) we identified lesion-outcome effects in men and women in the left-dominant language network, highlighting ventral fiber tracts of the left ILF, the IFOF, and the UF as core lesion focus across loss of different language facets. (2) Across both sexes, our analyses of brain lesion topographies pointed to an overarching structural organization reflecting the dual-stream concept of language and revealed pronounced sex divergencies in the role of lesion patterns. (3) We provide novel evidence for sex-nuanced lesion-outcome effects in the domain-general supra-modal networks, which our model attributed to cortico-subcortical pathways, with strong implications of the fornix and cingular fiber bundles. In the following, we discuss our findings for the anatomical topology of the identified lesion atoms, the shared lesion-outcome effects across sexes, and diverging predictive effects between men and women in implicated domain-general supra-modal networks.

Our analytical approach offered several advantages over classical lesion-symptom mapping studies. In conventional lesion-symptom mapping, each brain voxel is usually considered in isolation (Bates et al., 2003). However, analyzing the contribution of each single brain voxel individually deters intuitive interpretation by omitting interaction effects between spatially distant locations. Multivariate extensions have started to address the problem of mutual dependence between voxels (Pustina et al., 2018; Smith et al., 2013; Sperber, 2020). However, these approaches remain ill-suited to detecting overarching sources of variation that are driving disparate brain effects in continuous degrees, such as brain lateralization or sex-specific lesion-behavior associations. By means of our purpose-designed generative model, we could simultaneously appreciate local tract-lesion effects (and inter-dependencies between them) and their spatially distributed configurations. Leveraging the hierarchy of our fully probabilistic approach, we can directly integrate their hemispheric and sex-specific idiosyncrasies into one unified model for clinical outcome prediction.

The estimation of our framework was supported by empirical lesion data from a large sample of 1,401 stroke patients with rarely available whole-brain coverage. In line with the existing literature, the lesions were predominantly localized in subcortical zones and led to a critical disconnection of deep white matter tracts (Corbetta et al., 2015). Our data source thus presented two thrusts forward in stroke neuroimaging modeling. First, the spatial topography of the discovered lesion constellations covered both hemispheres. In many previous stroke studies on aphasia or spatial neglect, researchers have limited the anatomical space of investigation to either the left or right hemisphere alone (Baldo et al., 2013; Basilakos et al., 2015; Bates et al., 2003; Harvey & Schnur, 2015; Mirman et al., 2015; Smith et al., 2013). This unique data scenario enabled us to detect delicate predictive effects, including subtle sex differences, with regard to both brain hemispheres. Second, our findings provide comparable lesion coverage for both sexes. In contrast to the large majority of stroke lesion studies (Baldo et al., 2013; Mirman et al., 2015; Wu et al., 2015), we explicitly model sex as a higher model effect of interest. Potential differences independent of the sex-specific brain damage-behavior relation are further accommodated by our covariates at the lower level of our Bayesian model. This strategy ensures unique lesion-symptom effects for women and men and potentially helps to explain previous seemingly contradictory findings (Cahill, 2006).

Our pattern discovery framework enabled us to identify topologically coherent and biologically meaningful spatial patterns of lesion distributions from stroke. The identified lesion atoms recapitulated well-known functional brain circuits. Moreover, the respective lesion patterns generally fit well with the associated functions engaged in the applied tasks, as evidenced by neuropsychological assessments of language, motor, memory, and attention, as well as cognitive and executive control (Duncan & Owen, 2000; Fedorenko et al., 2013; Gregory Hickok & Poeppel, 2007). The fine-grained nature of the extracted lesion atoms also enabled us to bring into sharp focus fiber pathways of essential relevance for the human language capacity. The implicated lesioned anatomical tracts recapitulated the dorsal and ventral route of the dual-stream concept of language processing (Gregory Hickok & Poeppel, 2004, 2007), cerebral and cerebellar motor representations (Schmahmann et al., 2000), and visuo-spatial attention and executive control (De Schotten et al., 2011; Duncan & Owen, 2000). Overall, our analyses on language performance show a stronger impact of lesions in the left hemisphere, conforming to the widely held asymmetry of the human language network (Hartwigsen et al., 2021).

The granularity of our lesion atoms further enabled us to dissect subcomponents of stroke effects in language-related pathways, especially the different perisylvian subsegments of the AF. Since the three-segment view of AF anatomy was proposed (Catani et al., 2005), various studies ventured to examine the role that subcomponents of the AF play in realizing language function. Such studies initially suggested that the functions of the distinct fiber bundle segments differ: the direct pathway was associated with phonological processes, whereas the indirect anterior segment tended to serve semantically-based functions, and the posterior segment could serve especially language comprehension (Catani et al., 2005). However, evidence from these studies have previously been hard to reconcile.

Some studies on aphasia after stroke linked the anterior segment of the left AF to impaired speech fluency, among other fiber tracts (Basilakos et al., 2014; Fridriksson et al., 2013). One study advocated significant language associations selectively for the posterior segment of AF (Ivanova et al., 2016). Yet, other investigators reported no significant links between the damage to the subsegments of the AF and subsequent language function (Q. Yu et al., 2019). Based on our investigation of a large clinical cohort, we provide newly detailed evidence that dissects the major white-matter pathways necessary for language production and understanding. In particular, our findings highlight the anterior and long segments as key subcomponents of the perisylvian route. Within the lesion atom of the dorsal pathway, both the anterior and long segments of the AF are further confirmed in the prediction of speech fluency in men and women. Indeed, our lesion atoms reflecting the left and right ventral language route further include the posterior segment of the AF. This finding suggests a shared essential piece of the dorsal and ventral language pathway and a contribution of the latter pathway to naming and semantic fluency deficits. Our observation ties into a previous lesion study which reported overlap between both streams in the left temporo-parietal region (Fridriksson et al., 2016). Going beyond a merely topographic interpretation, our study extends the previous findings by pointing to potentially shared underlying anatomical connections. Such an overlap seems plausible given that the associated language tasks, naming, and semantic fluency, likely invoke both fiber processing streams.

In both men and women, the strongest observed brain-behavior predictions were preferentially lateralized to the left hemisphere. We identified the fiber tracts of the left ILF, the IFOF, and the UF to be critically involved in naming and semantic fluency deficits in both sexes. The left AF was responsible for performance decline in phonemic fluency in women and men to a similar degree, which nicely mirrors earlier findings (Marchina et al., 2011; J. Wang et al., 2013). The involved anatomical tracts are in accord with incumbent theories around the dorsal and ventral language pathways (Gregory Hickok & Poeppel, 2004, 2007; Saur et al., 2008). Our lesion-outcome effects in semantic fluency and naming were shared by women and men. This observation highlights the importance of the ventral language route and some of its major white matter pathways, including the ILF, IFOF, and UF. These findings directly converge with previous analyses that collapsed across both sexes into crude average effects. In particular, our finding linking semantic fluency with the ventral stream resonates well with vast evidence from large-scale neuroimaging meta-analyses (Binder et al., 2009; Vigneau et al., 2006), neurostimulation results in healthy participants (Pobric et al., 2007; Woollams, 2012), lesion studies (Bonilha et al., 2019; Fridriksson et al., 2016), computational modeling studies (Ueno et al., 2011), and targeted intraoperative electro-stimulation of fiber tracts (Duffau et al., 2005; Leclercq et al., 2010).

Our analyses detailed brain-behavior associations of phonemic fluency to locate to the dorsal perisylvian route, congruently for both sexes. With the strong impetus of the AF fiber tract, our results are consistent with the association of the dorsal language stream with phonological processing in healthy volunteers (Catani et al., 2005; Parker et al., 2005; Saur et al., 2008) and lesions in stroke patients (Fridriksson et al., 2016). Our predictive effects link the ventral and dorsal pathways to semantic and phonemic fluency, respectively. Both identified associations are aligned with findings from a variety of methodologies, including direct intraoperative electro-stimulation (Duffau et al., 2005; Leclercq et al., 2010), diffusion MRI tractography (Glasser & Rilling, 2008; Parker et al., 2005), and lesion-symptom investigations in aphasia (Biesbroek et al., 2016; Butler et al., 2014; Faroqi-Shah et al., 2014; Kümmerer et al., 2013; Mirman et al., 2015; J. Zhang et al., 2018). We confirm the structural relations in the dual-stream model in regard to the underlying language tasks. Our lesion findings shared by women and men are hence consistent with previous studies that were blind to sex-specific effects.

More generally, several previous efforts compounded evidence that male and female brains are distinct in the asymmetry of functional brain organization. Male brains appear to show more far-reaching segregation between the left and right hemispheric function. Instead, female brains are thought to have stronger interconnections between the left and right hemispheres (Jaeger et al., 1998; Shaywitz et al., 1995). According to an early hypothesis, the sharper distinction between left and right brain function, allegedly characteristic for men, was thought to promote spatial but impede verbal performance (Levy, 1978; Shaywitz et al., 1995). On a cognitive level, this sex divergence in brain lateralization was thought to explain why women outperformed the opposite sex in language tasks such as verbal fluency tests on average; as well as why men were supposed to typically exhibit greater spatial skills, such as in three-dimensional mental rotation tasks (Broverman et al., 1972; Hyde, 2016). However, the respective effect sizes have been small and several meta-analyses called into question the organizational differences related to lateralization and language processing (Hirnstein et al., 2019; Ruigrok et al., 2014; Wallentin, 2009).

In our sex-sensitive analyses, we found women to be more impaired by lesion damage in the left ventral pathway across different language tasks, including phonemic and semantic fluency. Moreover, we detailed a female-specific selective vulnerability to ventral lesion damage in phonemic fluency. Importantly, differences in lesion size or overall lesion topography were accounted for in our approach. A relatively stronger role of the left ventral pathway in women is in line with previous work in healthy volunteers, demonstrating higher radial diffusivity in the left UF in women relative to men (M. Jung et al., 2019).

In contrast, damage in the left-dominant ventral pathway and its right-hemispheric homolog was particularly informative about lost naming and semantic fluency capacities in men. Previous work suggested that the ventral language pathway is largely bilaterally organized, comprising parallel processing streams (Gregory Hickok & Poeppel, 2004, 2007). We expand the current literature body by adding novel evidence for a predominantly left-lateralized ventral dominance in women. In support of our findings, early analyses stratified by sex suggested that men have a symmetrical increase in white matter volume and concentration in the bilateral anterior temporal lobe (Good et al., 2001). Our findings receive further support in that healthy male volunteers showed increased grey matter volume and cortical thickness in several bilateral regions relative to females, although no relationship with speech tests was reported (Angelopoulou et al., 2019; Ritchie et al., 2018). Notably, previous studies are inconclusive with respect to sex incongruencies in functional and structural substrates of language-related processes (Ullman et al., 2007; Xu et al., 2020). Our sex-aware analyses bear the potential to reconcile several important inconsistencies on structural lateralization effects in general and in the ventral language pathway in particular.

Indeed, our analyses brought into the open several sex-specific findings outside the canonical language circuits that shed new light on stroke-induced language deficits. Our sex-aware analysis framework offered evidence that men draw from bilateral ventral and cerebellar connections and potentially integrate more domain-general cognitive and executive functions via right-lateralized cingular fiber connections to enable efficient language processing after stroke. There is increasing evidence that many higher cognitive functions, including language processing, are supported by large-scale distributed networks (Braga et al., 2020; Fedorenko et al., 2013). Such parallel organization of networks may potentially aid recovery after aphasic stroke by the recruitment of other cognitive functions that compensate for the damaged language capacities (Brownsett et al., 2014). In particular, additional domain-general networks may be upregulated to boost the performance of the damaged processes if the primary language system is compromised (Geranmayeh et al., 2014, 2017; Stockert et al., 2020). Damage to cingular fiber tracts may potentially undermine the increased recruitment of domain-general networks, including the ‘‘multiple-demand network’’ with special contributions to cognitive control (Duncan & Owen, 2000). The selective vulnerability we show for men in cingular pathways and bilateral temporal circuits may relate to previously identified sex-specific idiosyncracies in white matter microstructure in healthy subjects (Inano et al., 2011; Ritchie et al., 2018). The combination of core language network disruption and lesioned cingular pathways may hence reflect the complex balance of domain-specific and domain-general interplay for entertaining language functions.

In contrast, women accentuate left-dominant circuits, including structural connections of the fornix. A hippocampal-neocortical pathway through the fornix was previously suggested to be critical to the supra-modal brain network architecture of the default mode network (DMN, (Kernbach et al., 2018)). Large-scale associations networks, such as the DMN, fronto-parietal control, and salience networks, are generally thought to occupy regions juxtaposed and interdigitated with the language network (Braga et al., 2020; Margulies et al., 2016; Thomas Yeo et al., 2011). Altered supra-modal interactions with core language features via fornical fibers may hence reflect a dynamic process underlying neural remodeling in chronic aphasic stroke. Importantly, our findings capture the subacute phase of stroke; therefore the dynamic of reorganization may still change. Possibly, the altered language-DMN unit may affect language outcomes. Previous work has described disrupted DMN function in chronic stroke patients collapsed across sexes for impairment in motor function (Bonkhoff, Schirmer, Bretzner, Etherton, et al., 2021; Y. Zhang et al., 2017; Zhao et al., 2018) as well as language function (Balaev et al., 2016; Siegel et al., 2016; Tuladhar et al., 2013; C. Wang et al., 2014). Both the language-specific as well as domain-general disruption were suggested to influence post-stroke language outcome. However, the exact associations remain to be delineated.

Our findings speak in favor of revising the view towards sex-differentiated disruption of DMN integration. General sex divergences in DMN function in healthy individuals were previously demonstrated, indicating that women have higher connectivity within the DMN (Ritchie et al., 2018). The authors suggested that the higher intra-DMN connectivity may help explain the higher average female ability in different cognitive domains. The selective vulnerability to the fornix in women is further supported by a previous diffusion tensor investigation on healthy volunteers showing higher fractional anisotropy in women specifically in the hippocampus output pathway via the fornix (Inano et al., 2011). Moreover, in chronic stroke patients, the hippocampal-neocortical connectivity to the inferior parietal lobule (IPL) was consistently found to be decreased compared to healthy controls (J. Jung et al., 2021).

In fact, the IPL represents one potential hub for multi-sensory integration of domain-specific and supra-modal processes (Kernbach et al., 2018). Previous work has linked the IPL with semantic aspects of language processing, social cognition, and critical switch in stimulus-driven control of attention and diversion of self-reflective thinking to processing salient external stimuli (Bays et al., 2010; Binder & Desai, 2011; Bzdok et al., 2016; Kernbach et al., 2018). It receives axonal inputs from the posterior segment of the AF, as a short vertical tract connecting Wernicke’s with Geschwind’s areas (Catani et al., 2005). Our results further emphasize the relevance of the IPL and underlying posterior AF as seen in the dominance of the ventral lesion atom in women across all applied language tasks. Our findings converge with evidence suggesting negative modulatory connections from the parietal lobule to the superior temporal gyrus in females, enabling highly efficient semantic representations (Xu et al., 2020). Similarly, higher regional efficiency was reported in females in the left IPL and superior temporal regions in network connectivity analysis in neurologically intact participants (Gong et al., 2009).

Building on studies on healthy individuals, we confirm the relevance of the left IPL, fornix, and DMN in ischemic stroke patients. Our lesion-outcome findings in women may potentially reconcile previous observations and offer an explanation to sex divergences in chronic aphasia outcomes. More importantly, the personalized prediction of long-term post-stroke recovery using sex-aware stroke lesion topology may bring value to future individualized treatment interventions. Intensive speech therapy was previously shown to promote more efficient DMN integration leading to consequent language improvement (Abo et al., 2004; Dreyer et al., 2021; Marcotte et al., 2013; Musso et al., 1999). Targeted rehabilitation in men and women may hence advance therapy-induced neuroplasticity by enabling more efficient multimodal network integration.

## Online Methods

### Prospective stroke population resource

We analyzed a multi-center stroke registry with 1,401 patients. They were retrospectively selected from the Bundang and Hallym Vascular Cognitive Impairment cohorts, which are prospectively recruited cohorts consisting of patients originally admitted to the Seoul National University Bundang Hospital or Hallym University Sacred Heart Hospital in South Korea between 2007 and 2018 (Kim et al., 2015). Patients were hospitalized and diagnosed with acute ischemic stroke. All patients were eligible for the present study based on the following criteria: (1) availability of brain MRI showing acute tissue infarction in the diffusion-weighted imaging (DWI) and/or fluid-attenuated inversion recovery (FLAIR), (2) successful lesion segmentation and registration, (3) no primary intracerebral hemorrhage, and (4) availability of follow-up data on key demographics and neuropsychological assessment (the 60-min Korean-Vascular Cognitive Impairment Harmonization Standards-Neuropsychology Protocol (Hachinski et al., 2006; K.-H. Yu et al., 2013)). The local institutional review boards of each hospital approved the study protocol and gave waived consent requirements based on the retrospective nature of this study and minimal risks to participants.

### Characteristics of stroke patient cohort

The neuropsychological performance of the patients was profiled by a rich battery of tests ∼3 months (Hallym: 91.4 days (SD 38.9), Bundang: 105.7 days (SD 94.1)) after the acute onset of stroke (K.-H. Yu et al., 2013). For the purpose of our present study, we primarily focused on two key facets of post-stroke language performance: i) naming ability (Korean short version of the Boston Naming Test (BN) (Kaplan et al., 1983)), and ii) speech fluency (phonemic and semantic fluency (PF, and SF, respectively; (K.-H. Yu et al., 2013)). The BN task prompts patients to name 15 pictured nouns (short version). In the semantic fluency tasks, patients are instructed to generate as many words as possible belonging to a specific category, such as animals, within a one minute time window. In the phonemic fluency task, in turn, patients are asked to generate words starting with a given phoneme (e.g., giut, iung, or siut, which correspond to the international phonemic alphabet phonemes of g, ŋ, and s, respectively). These fluency tests have been previously validated in the Korean language (Kang et al., 2000). A diverse collection of general cognitive tests included the Korean Mini-Mental State Examination (MMSE), the Seoul Verbal Learning test (SVLT), the Rey Complex Figures test (RCFT), Digital symbol coding, and the Korean-Trail Making Tests Elderly’s version A and B (TMT). The Informant Questionnaire on Cognitive Decline in the Elderly (IQCODE) captured the cognitive performance before the ischemic event (D. W. Lee et al., 2005). Available sociodemographic and clinical information included age, sex, years of education, and lesion volume (**Table 1)**. All continuous variables were adjusted to a comparable scale by mean centering to zero and variance scaling to one.

### Neuroimaging data preprocessing

Whole-brain MRI scans were typically acquired within the first week after the stroke event. The scanning protocol included structural axial T1, T2-weighted spin-echo, FLAIR, and DWI sequences (3.0T, Achieva scanner, Philips Healthcare, Netherlands; cf., supplementary materials for details). Tissue damage was manually segmented on DWI or less frequently FLAIR images by experienced investigators (A.K.K. and G.A.) relying on in-house developed software based on MeVisLab (MeVis Medical Solutions AG, Bremen, Germany; (Ritter et al., 2011)). Lesion segmentations were checked and potentially refined by two additional experienced raters. Brain scans and their corresponding lesion segmentations were linearly and nonlinearly normalized to Montreal Neurological Institute (MNI-152) reference space employing the publicly available RegLSM image processing pipeline (Weaver et al., 2019). Quality of normalization was rigorously controlled by an experienced rater and if necessary manually corrected.

### Data-led deconvolution of hidden lesion atoms

We sought to explore the coherent patterns that may combine to make up the topographical distribution of stroke lesions in white matter. First, we parsed each patient’s lesion fingerprint by summarizing the white-matter damage annotations at the voxel level according to fiber tract definitions from a reference tractography atlas (Catani & Thiebaut de Schotten, 2008; Thiebaut de Schotten et al., 2011). This widely-used atlas provided spatial definitions of 28 major fiber tracts in MNI standard space (Assaneo et al., 2019; Corbetta et al., 2015; Fridriksson et al., 2013; Lunven et al., 2015; Zeestraten et al., 2017). In doing so, the tract-by-tract summaries of lesion loads separately tracked the left and right hemispheres, which enabled the modeling of subtle differences in hemispheric lateralization in downstream analysis steps (Catani et al., 2007; Thiebaut de Schotten et al., 2011).

Of particular relevance to the present investigation due to their roles in human-defining language capacity, we captured the neuroanatomy of the core perisylvian circuitry by the canonical arcuate fasciculus (AF), subdivided into i) long, ii) anterior, and iii) posterior segments of the tract (Catani et al., 2005; Catani & Mesulam, 2008; Catani & Thiebaut de Schotten, 2008; Dick & Tremblay, 2012). The ventral pathway consisted of the uncinate fasciculus (UF), the inferior longitudinal fasciculus (IFL), and the inferior fronto-occipital fasciculus (IFOF) (Axer et al., 2013; Catani et al., 2003; Saur et al., 2008; Vigneau et al., 2006). Limbic pathways included the fornix and cingular white-matter bundles (Catani & Thiebaut de Schotten, 2008). Motor projections were taken into account by the internal capsule (IC), the corticospinal tract (CST), and cerebellar connections, including the cortico-pontine-cerebellar tract (CPC) and the inferior and superior cerebellar peduncle (ICP, SCP, respectively). The topography of atlas fiber tracts directly informed the labeling of the derived lesion atoms. For consistency, the resulting lesion atoms of the language system were labeled according to widely acknowledged definitions of the dual-stream model of language (Gregory Hickok & Poeppel, 2004). We log_2_-transformed and concatenated the measures of each of the 28 tract measures of lesion loads for all 1,401 patients. Once each tract lesion load (Marchina et al., 2011; J. Wang et al., 2013) was recorded in each patient, we carried out non-negative matrix factorization (NNMF) as a multivariate pattern discovery strategy (D. D. Lee & Seung, 1999). This unsupervised machine-learning algorithm can identify the topological form and patient-specific combination of lesion pattern expressions that make up the white matter disconnection inflicted by stroke. In accordance with our recent stroke studies (Bonkhoff, Lim, Bae, Weaver, Kuijf, et al., 2021; Bonkhoff, Schirmer, Bretzner, Hong, et al., 2021), the derived sum-of-parts representation was henceforth referred to as *lesion atoms*.

More formally, NNMF achieves a low-rank approximation of the data *V*, with *V* reflecting the lesion load summaries of 28 tracts, with dimensions *m* x *n* (*m* = number of tracts in the atlas, *n* = number of patients), by partitioning the interindividual variation in white-matter strokes into a matrix *W* of *k* part-based factor representations. The latent pattern expression matrix *H* indicated how relevant each emerging lesion atom is to describe the constituent parts of an individual patient’s overall brain lesion distribution. *W* and *H* thus carried *m* x *k* and *k* x *n* dimensions, respectively. Given by *V = WH,* the latent factorization hence deconvolved the actual lesion constellation in a particular patient into a parts-based representation.

In contrast to alternative dimensionality reduction tools, NNMF provided at least two important advantages for the goal of the present study. First, the non-negative value encoding of the differential white-matter involvement and the inherent non-negativity constraint of the NNMF algorithm allows for intuitive neurobiologically meaningful interpretation. For each patient *j*, the latent pattern expression *H_j_* offered a directly interpretable model of the lesion variation across patients in our cohort. In contrast, alternative matrix factorization algorithms, such as principal component analysis, typically recover patterns by incurring various cancellations between positive and negative pattern loadings. Such alternative treatment of the derived hidden parts-of-whole representations would have hurt intuitive neurobiological meaning. By means of strictly additive combinations of pattern contributions (allowing for no subtractions), NNMF allows for biologically more meaningful and interpretable representations of lesion constellations. Second, classical clustering approaches for dimensionality reduction would consider the effect of each differential tract involvement once only. In contrast, by embracing NNMF, damage to each white-matter tract in the reference atlas has the opportunity to belong to multiple latent factors of *W* in continuous degrees. In this way, each tract lesion load could play a distinct role in different lesion atoms, each of which reflected extracted lesion-specific topological motifs distributed across the whole white matter.

### Predicting language outcome at 3 months post-stroke from patient-specific lesion fingerprints

The thus derived lesion atoms served as a low-dimensional summary of the individual lesion fingerprints. Expression degrees of these lesion patterns, specific to each particular stroke patient, provided the input to our Bayesian model (Gelman & Hill, 2006) to explain the interindividual differences in language performance outcomes at ∼3 months after the stroke event. We designed a Bayesian generative modeling tactic. We aimed to obtain fully probabilistic parameter estimates that could inform us about effect strength and effect certainty of how each particular lesion pattern is responsible for the clinical outcome. To place a special focus on the putative effects of sex divergences on post-stroke language performance, we introduced dedicated model parameters, which allowed to estimate each pattern regression slope in a sex-specific manner. That is, we could estimate the overlapping and distinct roles of each lesion atom in tracking cognitive facets of language in women and men separately within an integrated modeling solution. The model structure was as follows:

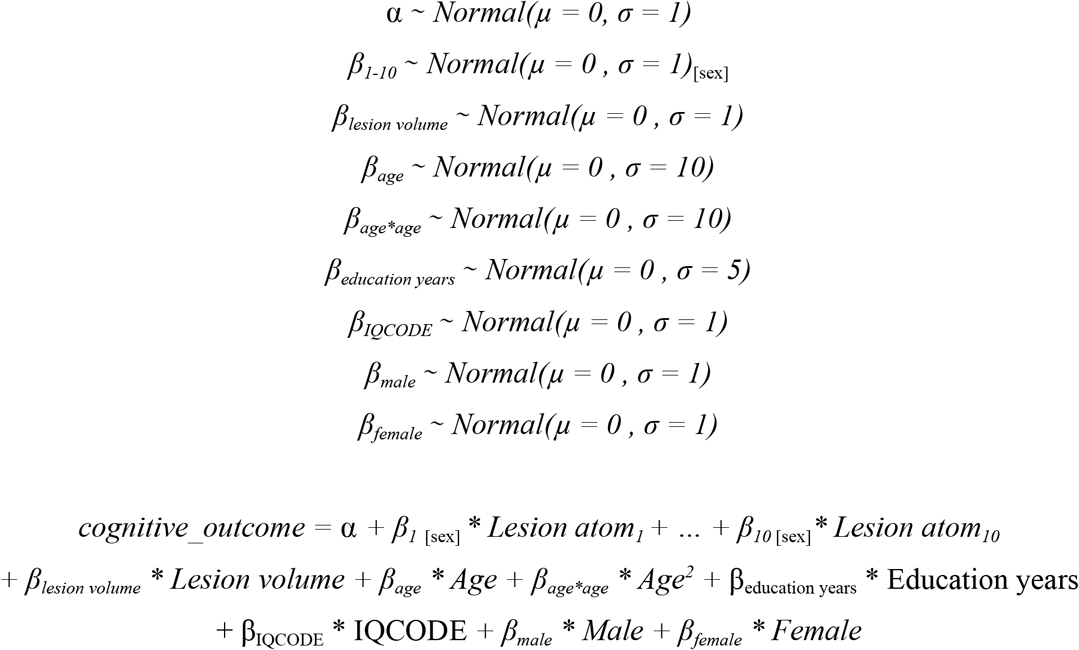

A separate probabilistic Bayesian model was estimated for each dedicated post-stroke performance score in each of the target language domains: BN, SF, and PF (cf. above; for other cognitive assessments see Supplement). All analyses were appropriately adjusted for variation that could be explained by the covariates age, age^2^, sex, education in years, pre-morbid cognitive performance, and total lesion volume. We adjusted for age-specific effects by considering the bare age and its squared value and, therefore, captured possible u-shaped effects. Our parametric model specification also included sex as a covariate to take into account outcome differences that were independent and dependent of the lesion configurations captured in the NNMF-derived lesion atom representations (see (Bonkhoff, Lim, Bae, Weaver, Kuijf, et al., 2021; Bonkhoff, Schirmer, Bretzner, Hong, et al., 2021)). Samples from the posterior distribution of the model parameters were drawn by the No U-Turn Sampler (NUTS, (Hoffman et al., 2014)), a kind of Monte Carlo Markov Chain algorithm (in our setting defaulted to draws=5,000).

To evaluate the practical usefulness of the thus inferred posterior parameter distributions, we computed posterior predictive checks for every analysis of a given language performance outcome. These model-based simulations of new data were then compared to the observed data to determine the predictive performance measured as coefficient of determination (R^2^). That is, we assessed the model-simulated outcome predictions generated by our Bayesian model to approximate an external validation based on our patient cohort. This empirical procedure is a well-recognized option for judging the adequacy of Bayesian models given the actual data at hand (Gelman et al., 2013; Kruschke, 2014).

### Code availability

Data analyses were conducted in a Python 3.8.5 (IPython 7.21.0) environment and primarily relied on the packages nilearn, scikit-learn, and pymc3 (Abraham et al., 2014; Salvatier et al., 2016). Full code is accessible to and open for reuse by the reader here: http://github.com/jkernbach/to_be_added_later. The probabilistic tractography atlas is available at: https://www.natbrainlab.co.uk/atlas-maps. The use of dynamic reporting guarantees full reproducibility of the results.

## Acknowledgments & Disclosure

We confirm that we have read the Journal’s position on issues involved in ethical publication and affirm that this report is consistent with those guidelines.

We gratefully acknowledge Angelina K. Kancheva and Gözdem Arikan for their help with performing the manual infarct segmentations, Jae-seol Park, and Eunbin Ko for their help in organizing the neuropsychological data, and Parashkev Nachev and Michel Thiebaut de Schotten for their valuable comments on earlier versions of the manuscript.

## Availability of data and code

Additional data is made available in the supplemental methods and supplemental results online.

## Supplementary Online Material

**Supp. Figure 1.**
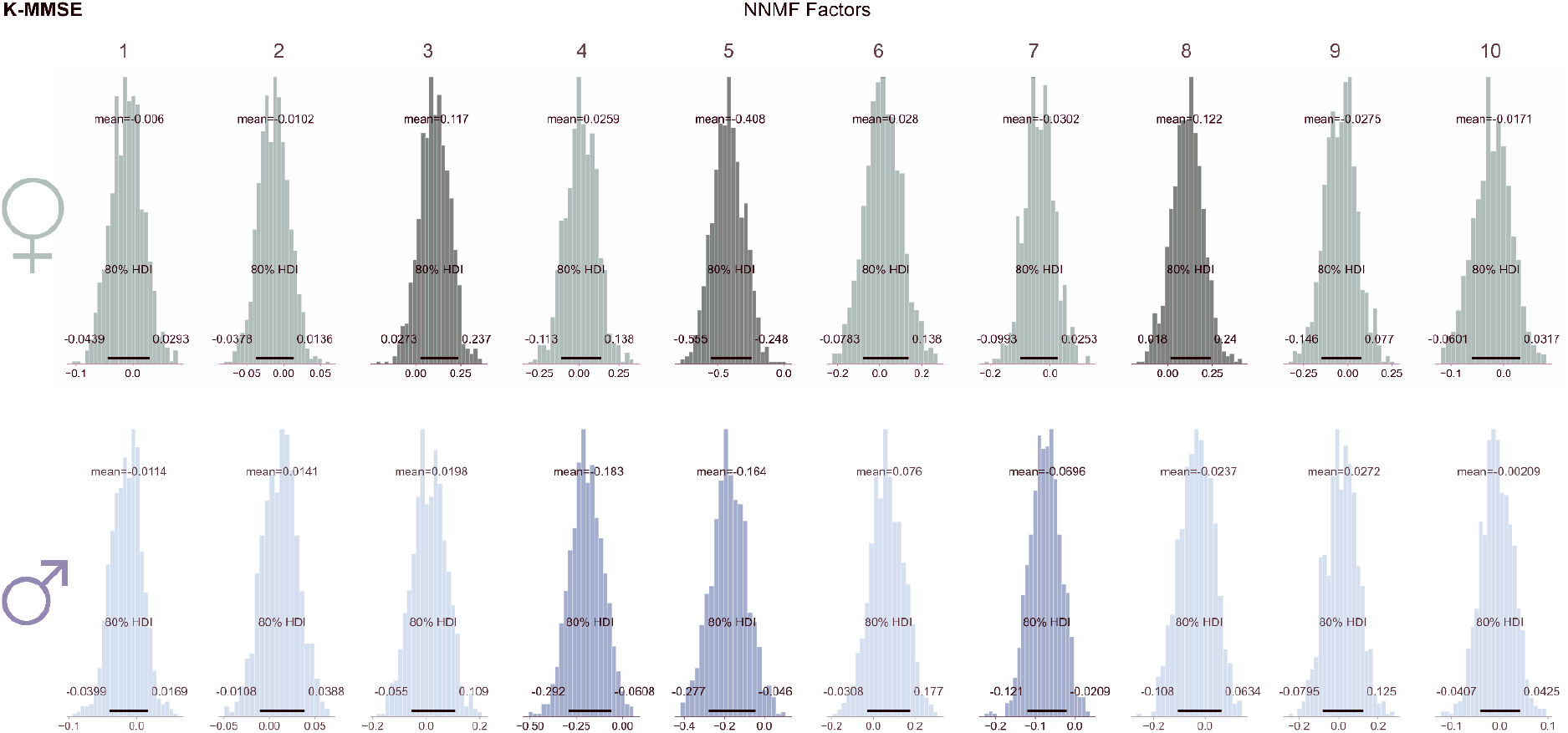
K-MMSE Bayesian posteriors.

**Supp. Figure 2.**
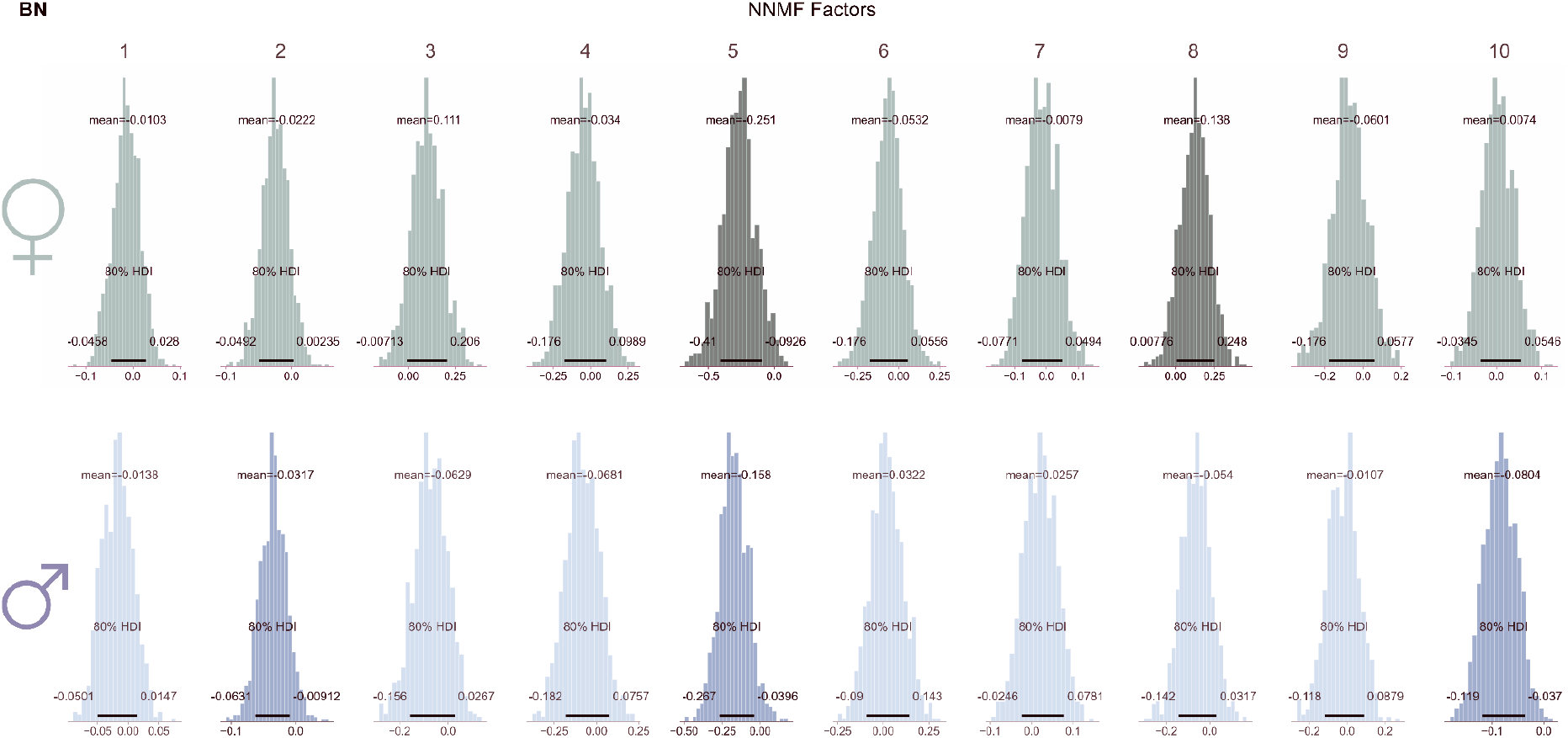
BN Bayesian posteriors.

**Supp. Figure 3.**
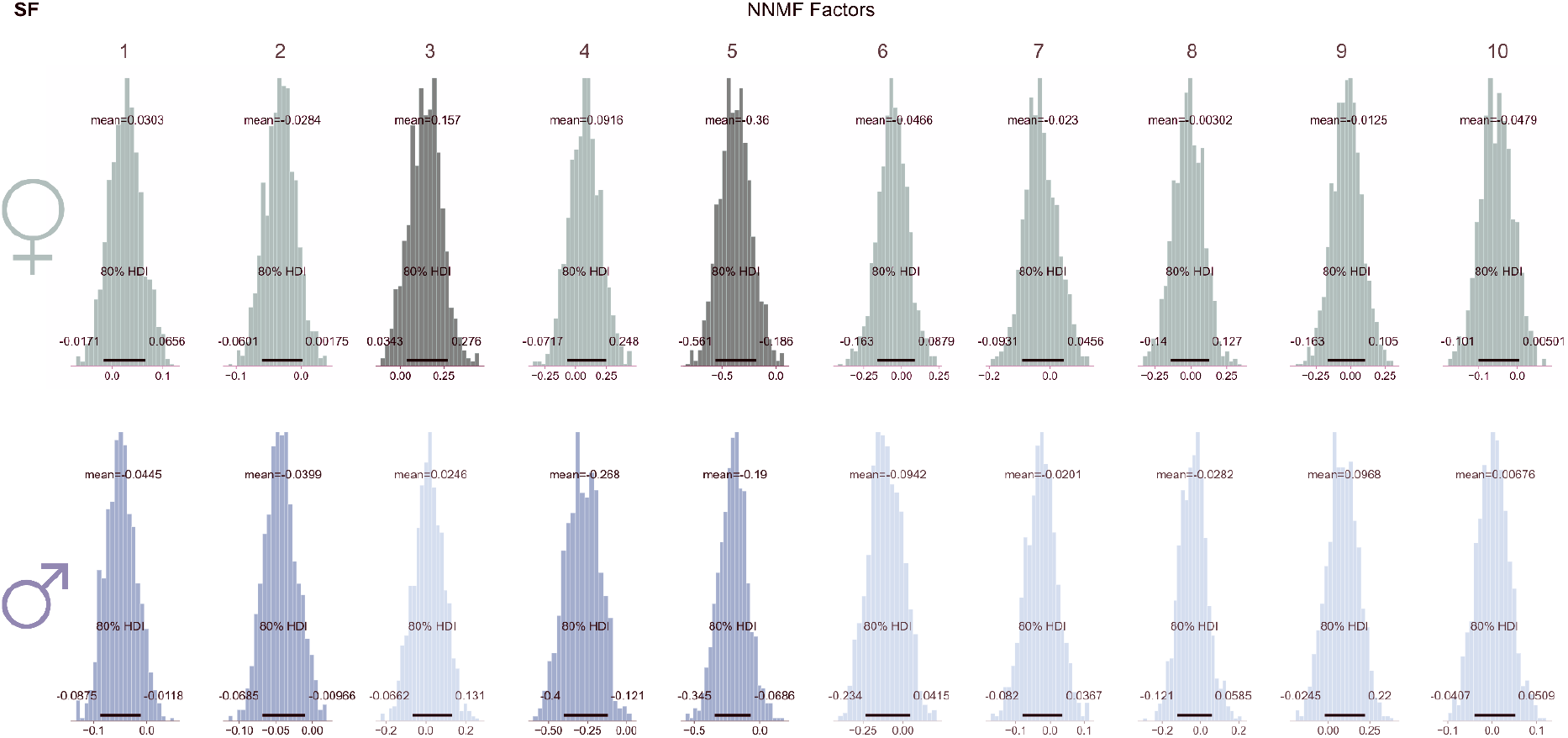
SF Bayesian posteriors.

**Supp. Figure 4.**
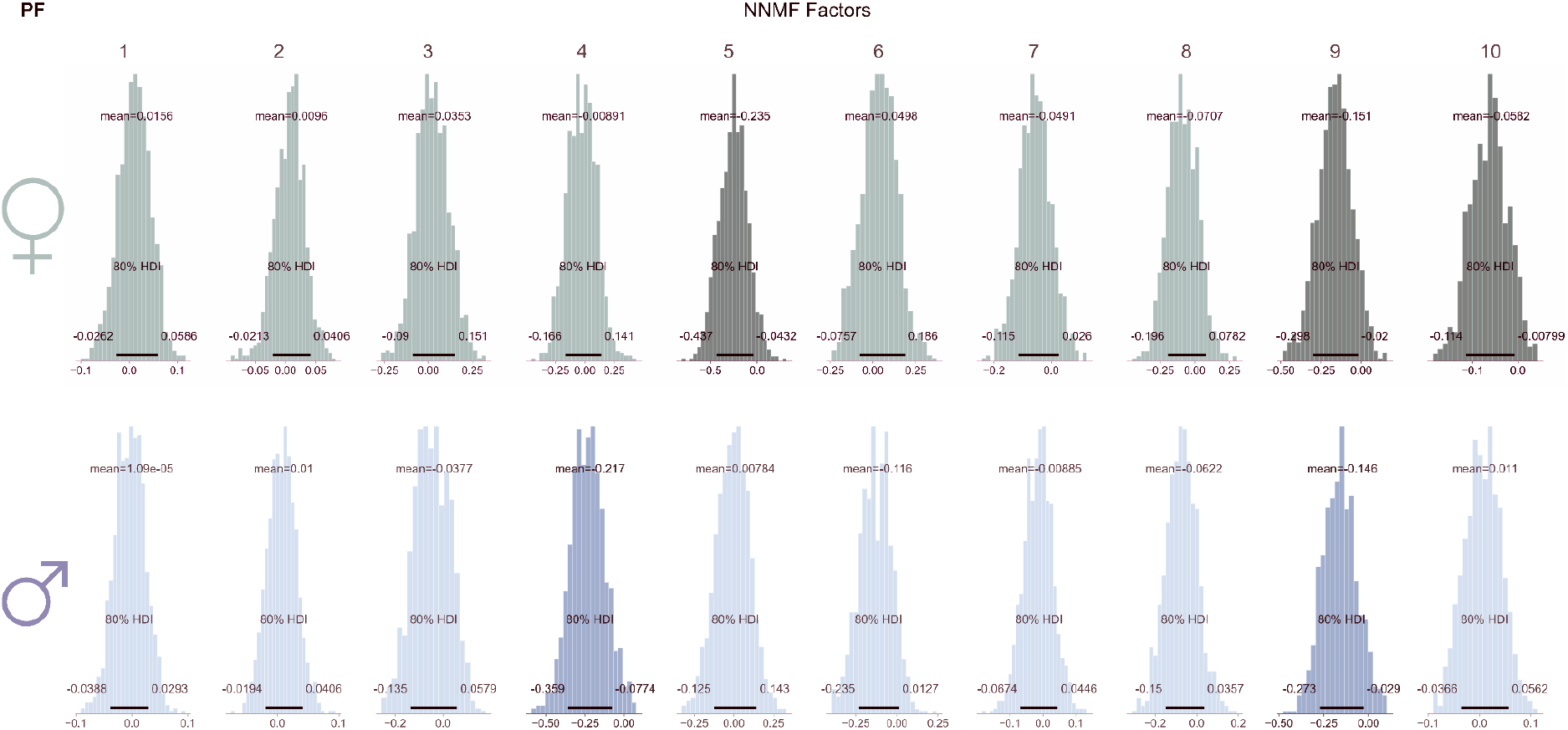
PF Bayesian posteriors.

**Supp. Figure 5.**
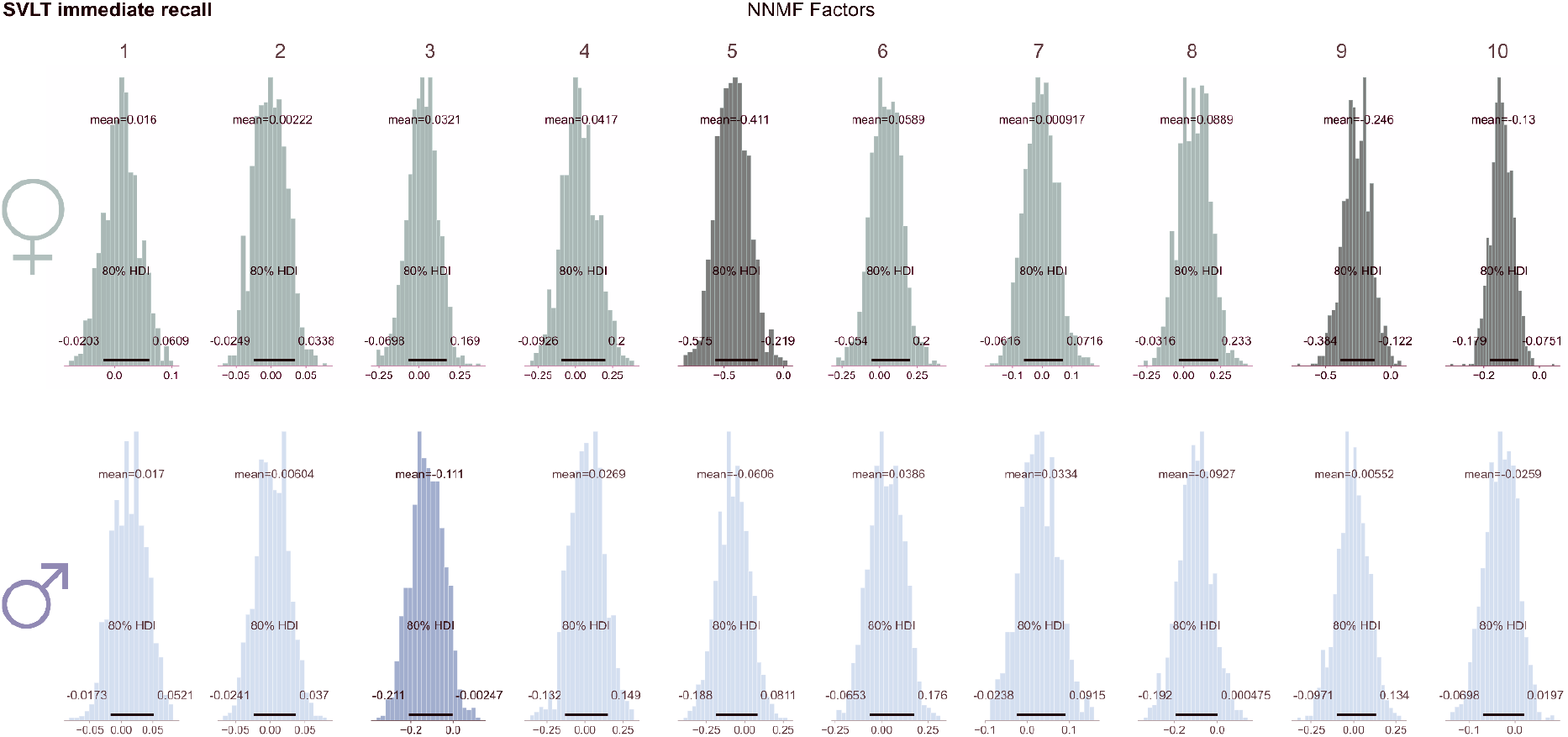
SVLT immediate recall Bayesian posteriors.

**Supp. Figure 6.**
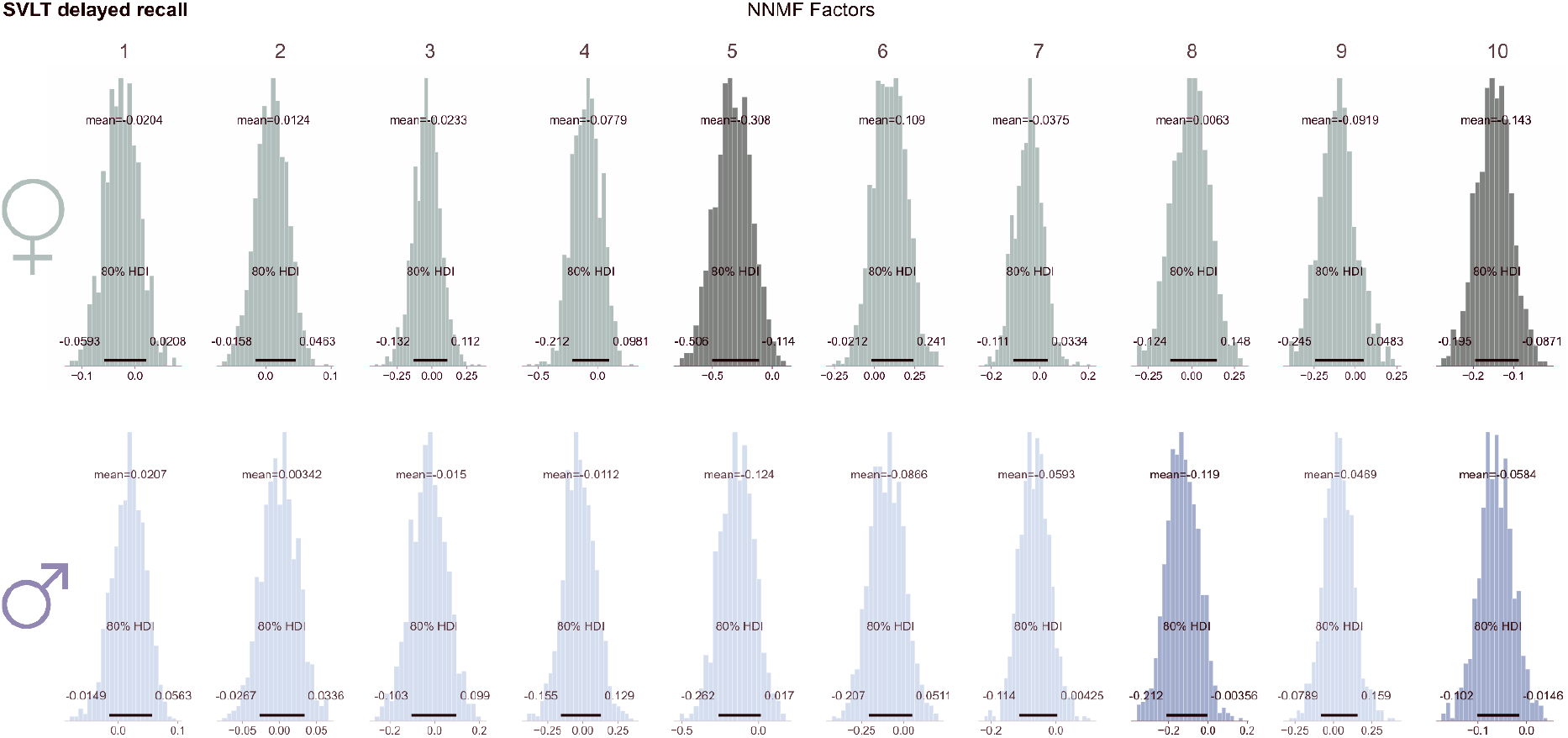
SVLT delayed recall Bayesian posteriors.

**Supp. Figure 7.**
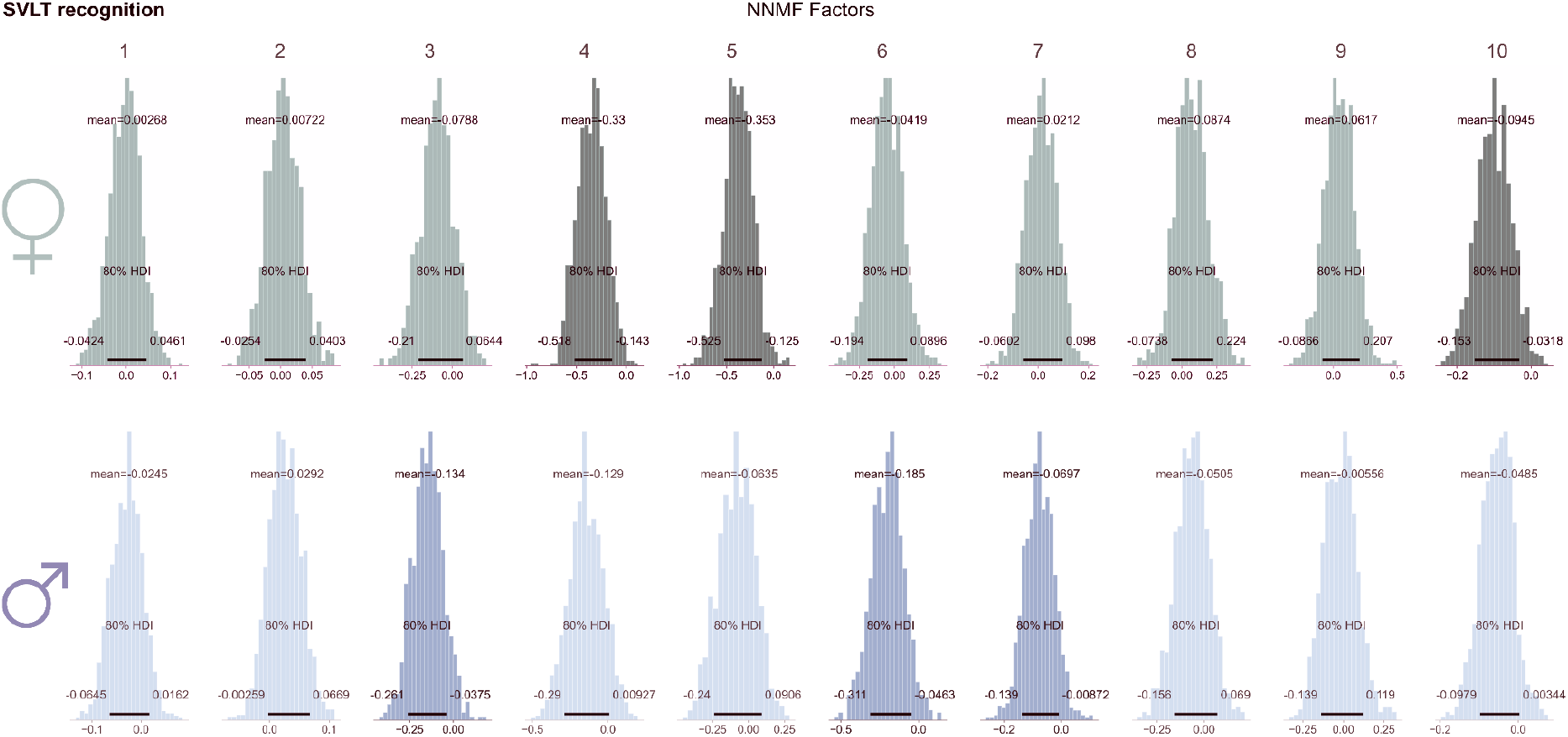
SVLT recognition Bayesian posteriors.

**Supp. Figure 8.**
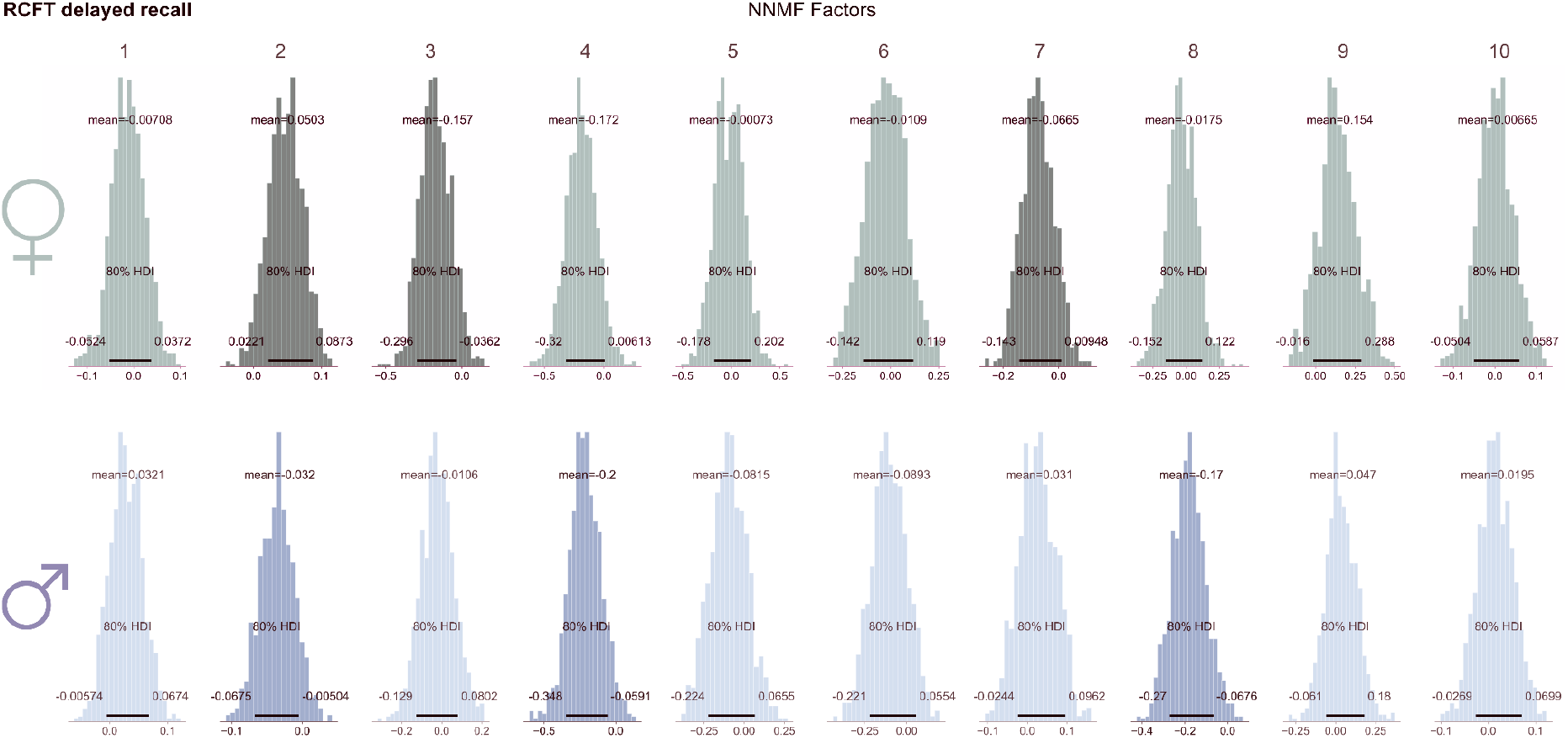
RCFT delayed recall Bayesian posteriors.

**Supp. Figure 9.**
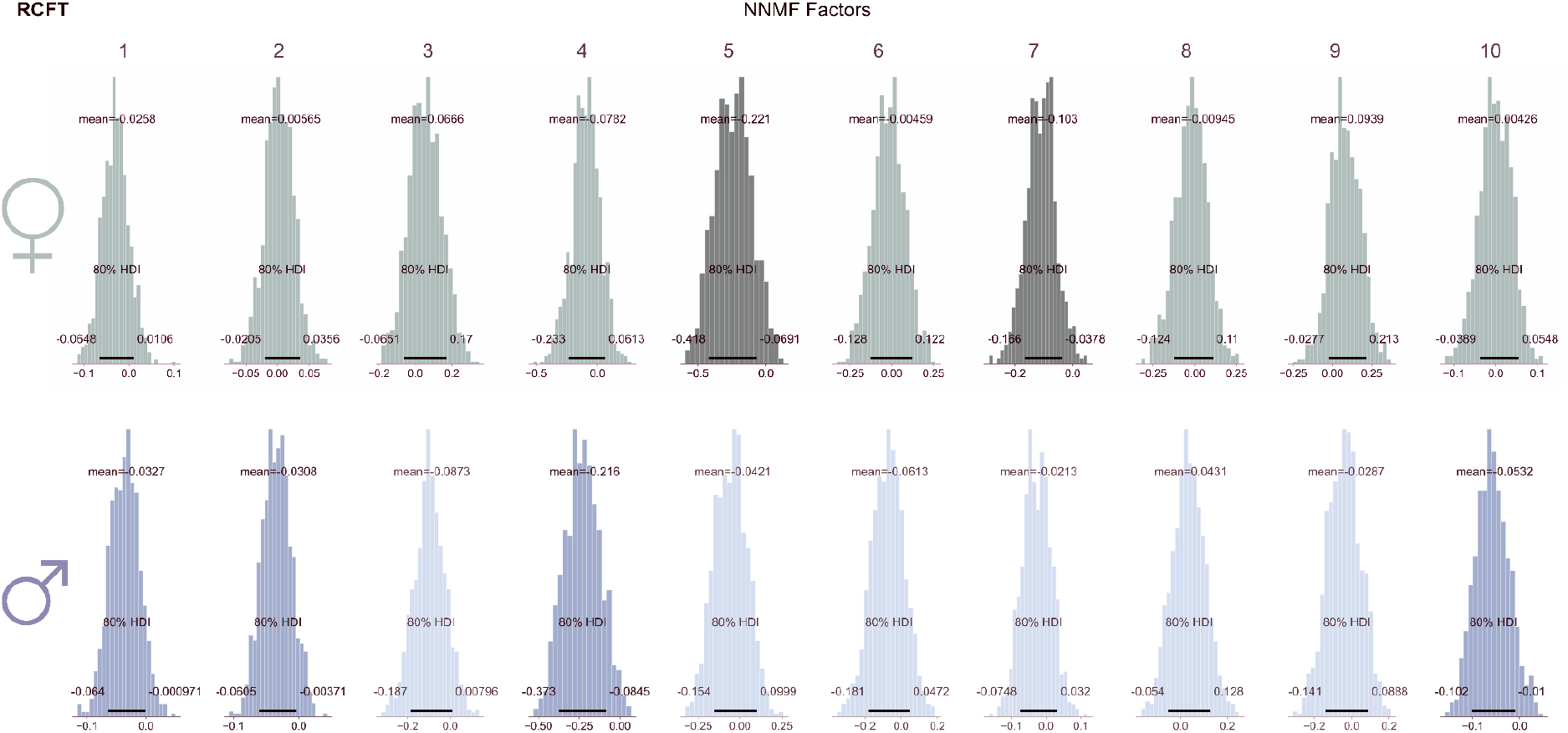
RCFT Bayesian posteriors.

**Supp. Figure 10.**
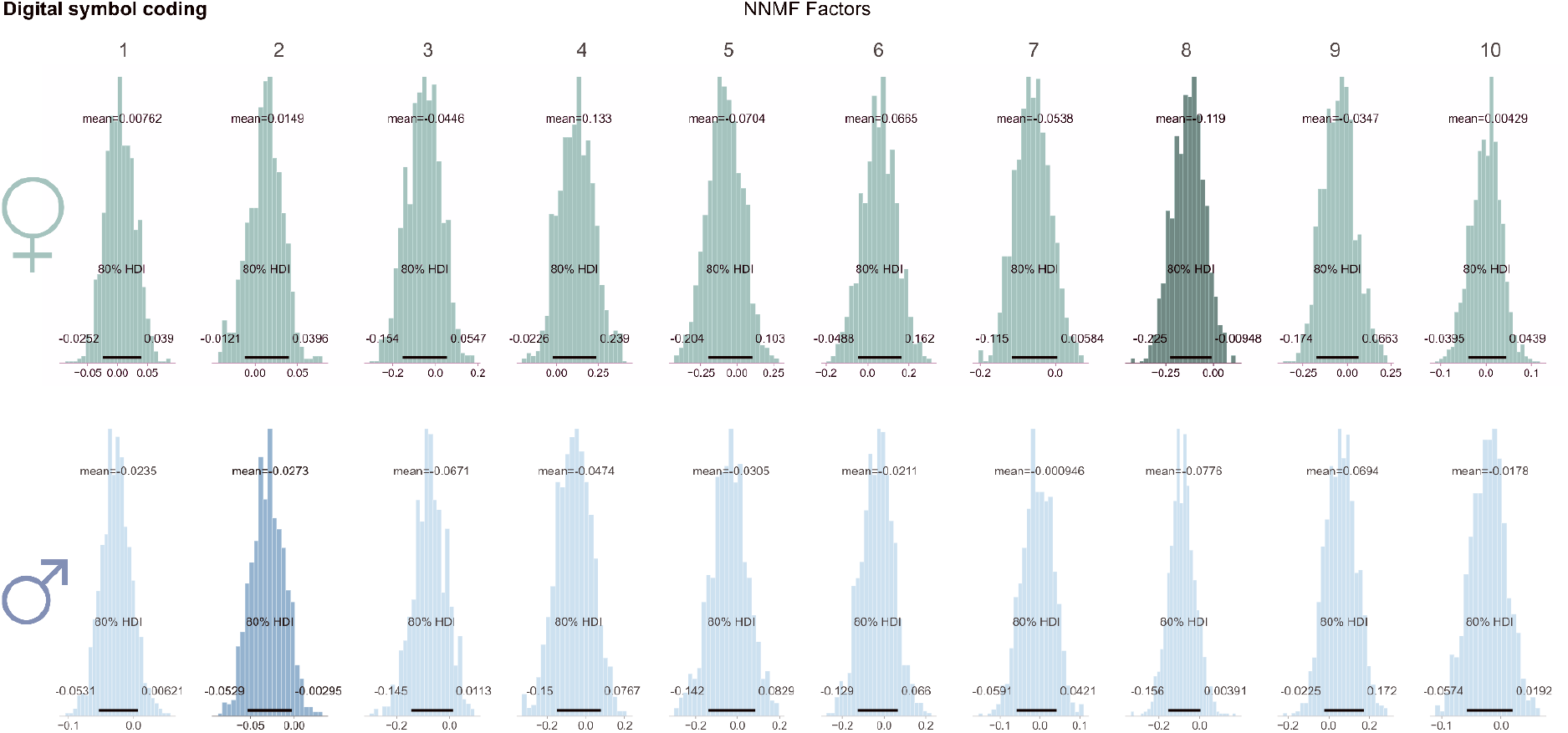
Digital symbol coding Bayesian posteriors.

**Supp. Figure 11.**
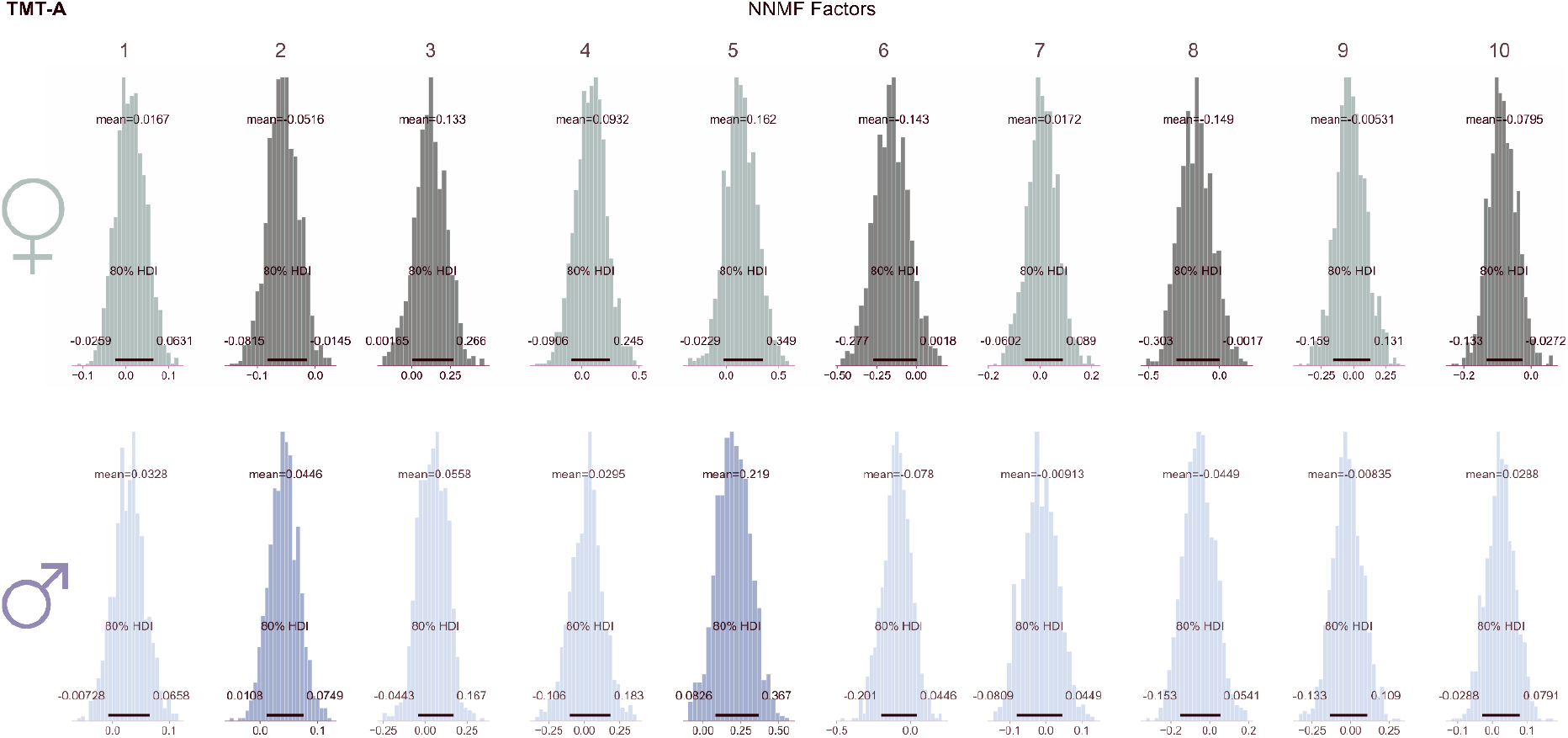
TMT-A Bayesian posteriors.

**Supp. Figure 12.**
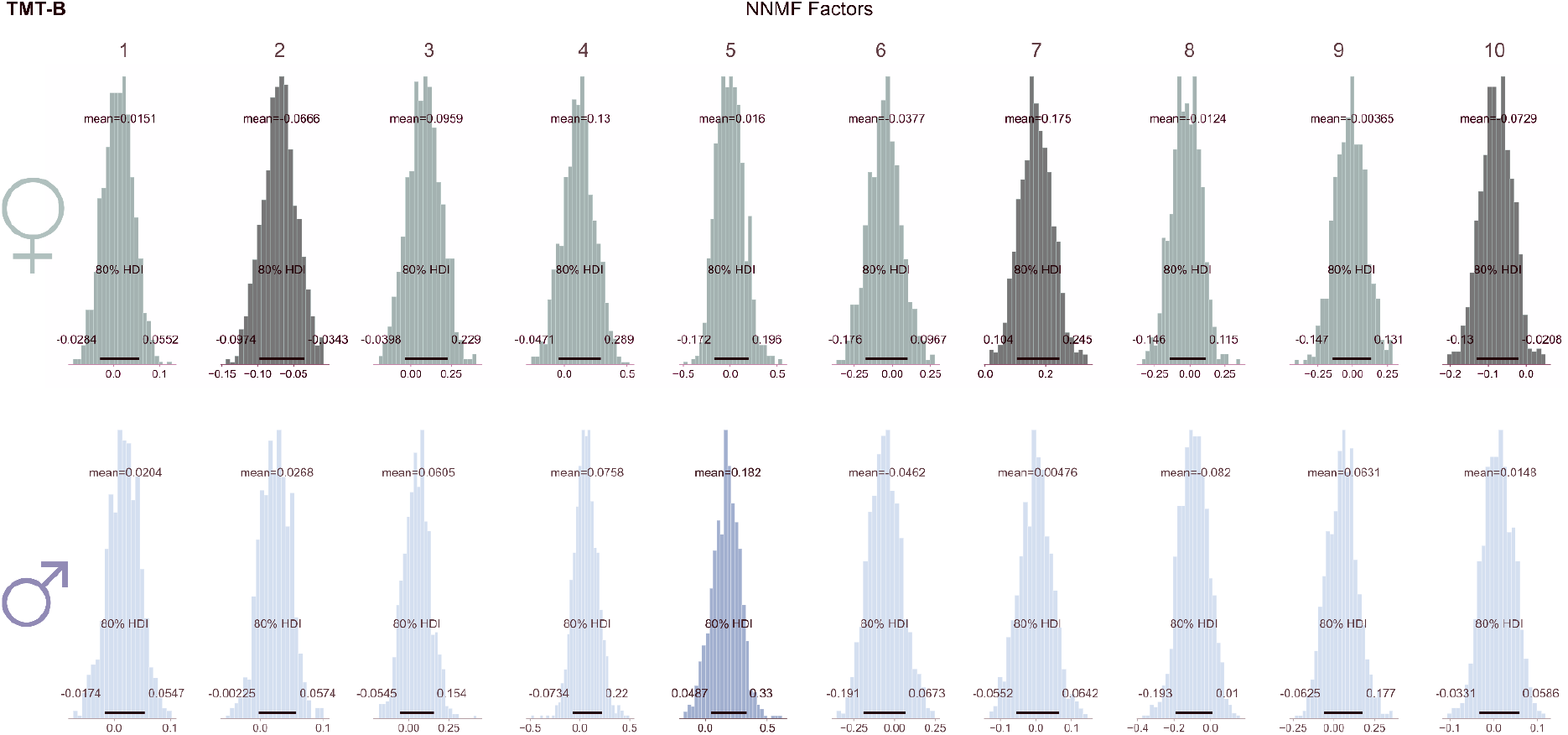
TMT-B Bayesian posteriors.

